# The evolution of age-specific resistance to infectious disease

**DOI:** 10.1101/2022.10.05.511061

**Authors:** Lydia J. Buckingham, Emily L. Bruns, Ben Ashby

## Abstract

Innate, infection-preventing resistance often varies between host life-stages. Juveniles are more resistant than adults in some species, whereas the opposite pattern is true in others. This variation cannot always be explained by prior exposure or physiological constraints and so it has been hypothesised that trade-offs with other life-history traits may be involved. However, little is known about how trade-offs between various life-history traits and resistance at different life-stages affect the evolution of age-specific resistance. Here, we use a mathematical model to explore how trade-offs with natural mortality, reproduction and maturation combine to affect the evolution of resistance at different life-stages. Our results show that certain combinations of trade-offs have substantial effects on whether adults or juveniles are more resistant, with trade-offs between juvenile resistance and adult reproduction inherently more costly than trade-offs involving maturation or mortality (all else being equal), resulting in consistent evolution of lower resistance at the juvenile stage even when infection causes a lifelong fecundity reduction. Our model demonstrates how the differences between patterns of age-structured resistance seen in nature may be explained by variation in the trade-offs involved and our results suggest conditions under which trade-offs tend to select for lower resistance in juveniles than adults.

## Introduction

Immunity to infectious diseases typically varies across the lifespan of the host, which has significant consequences for host health and disease transmission [1–3]. For instance, if the disease is more severe in adults than in children, then a population with high adult and low juvenile resistance will have fewer severe cases than a population which is homogeneous in resistance. Similarly, if adults and children have more contact with individuals in the same age class, then age-specific resistance will impact on the size of the epidemic.

Variation in different types of immunity (e.g. innate resistance, adaptive immunity, infection-preventing resistance, tolerance) with host age has been observed in many taxa, such as plants [4–7], invertebrates [8–12] and vertebrates [13–15], including humans [16–18]. Yet the nature of age-specific immunity varies widely, with adults better protected than juveniles in many [5–10,12–17] but not all cases [4,11,12,18]. Differences in age-related patterns of host immunity exist both within and between species [19–22], but the reasons behind these diverse patterns are not always well understood. In particular, we lack a detailed understanding of the factors affecting differences in selection for host defences against infectious disease at different host life-stages.

Variation in host immunity with age may occur due to a variety of mechanisms, including immune priming [23,24]; adaptive immunity [25,26]; the loss of maternal antibodies in mammals [27,28]; senescence [29,30]; the accumulation of pathogenesis-related (PR) proteins and activation of the salicylic acid pathway in plants [6]; dilution of pathogen effects due to changes in body size in insects [8]; differences in transcriptional responses to infection in molluscs [12] and changes in the ratio of naïve to memory T-cells in humans [16]. However, in many cases, the mechanisms which cause differences in juvenile and adult immunity are unknown or poorly understood [4,5,7,9–11,13–15,17,18]. When immunity depends on prior exposure, juveniles may be less resistant to infection simply because they have yet to experience pathogens that adults have previously encountered (although juveniles may be more resistant to infection than adults if immunity wanes over time). Whilst variation in prior exposure can contribute to patterns of age-specific immunity, especially in vertebrates, it cannot fully explain observed differences in juvenile and adult immunity. Such differences also exist, for instance, in species which rely solely or primarily on innate, rather than acquired, immunity [5–10,12] and when a population encounters a novel pathogen to which neither adults nor juveniles have acquired immunity [31].

From an evolutionary perspective, one might expect that innate (non-adaptive) defences against infectious diseases should always be greater in juveniles than in adults, since infection at a young age could lead to death or sterilisation before reproduction can occur [32]. However, this is not always the case. Trivially, physiological constraints may constrain juvenile defences in some species, preventing juveniles from evolving stronger protection against parasitism or herbivory [33,34]. Although this may provide a partial explanation for supressed juvenile defences, artificial selection for increased innate immunity [35–37] and evidence of polymorphism in the level of immunity in natural populations [19–22,38–40] have shown that many hosts do not possess the maximum possible level of juvenile immunity. Hence physiological constraints on juvenile defences do not provide a full explanation. Differences in disease outcomes may also drive selection for age-specific immunity, for example if a disease causes higher virulence in adults than in juveniles (see appendix D of [41]). Again, this may be a factor in driving differences in immunity, but adult immunity has been found to be higher than juvenile immunity in systems in which the disease has the same effect on susceptible hosts of all life-stages [7]. Therefore, this cannot provide a complete explanation either.

An alternative evolutionary explanation for differences in juvenile and adult immunity is that host defences trade off with other life-history traits. For example, increased juvenile immunity may require resource allocation away from growth and development, resulting in a negative relationship between juvenile immunity and maturation, mortality or future reproduction. Similarly, adult immunity may require resources to be diverted away from reproduction or may be associated with higher mortality from other causes. There is empirical evidence for trade-offs between reproduction [42,43] or growth [42–45] and host immunity in plants and invertebrates, though little data is available on age-specific effects. The impact of these different trade-offs on the evolution of immunity across the host lifespan has yet to be determined theoretically.

To date, theoretical models have explored the spread of disease in age-structured populations [3] or the evolution of immunity in populations with no age structure [46–50]. However, the evolution of innate, infection-preventing resistance at different life-stages has received little attention. As an exception, Ashby & Bruns [51] explored the evolution of (innate) juvenile susceptibility to infection in a population with fixed adult susceptibility, under the assumption that juveniles are always at least as susceptible as adults. They found that juveniles may evolve higher susceptibility than adults under a wide range of conditions, but the difference was most extreme when hosts had very long or very short lifespans. Here, we build on these findings by allowing juvenile and adult resistance to evolve simultaneously and independently and by exploring how a range of trade-offs with different life-history traits affect the evolution of resistance across the host lifespan. As in Ashby & Bruns’ paper [51], we consider the specific case where resistance prevents infection (as opposed to resistance which limits or eliminates infection). We focus our analysis on trade-offs with maturation, mortality and reproduction, along with variation in pathogen traits, specifically transmissibility and the strength and type of virulence. We show that juvenile resistance is most costly when it trades off with reproduction later in life, resulting in lower juvenile resistance than evolves under other trade-offs and also lower juvenile than adult resistance (assuming equal strength of trade-offs). Furthermore, we show that a trade-off between juvenile resistance and reproduction can cause juvenile resistance to be lower than adult resistance even when infection causes a permanent reduction in fecundity.

## Methods

### Model description

We expand the model described by Ashby and Bruns [51] to explore the evolution of innate, infection-preventing resistance at juvenile (*J*) and adult (*A*) stages, in a well-mixed, asexual host population (see Fig. 1a for a model schematic and Table 1 for a full list of parameters and variables). Let *S_i_* and *I_i_* be the densities of susceptible and infected hosts respectively at life-stage *i* ∈ {*J*, *A*}, giving a total host population density of *N* = *S_J_* + *S_A_* + *I_J_* + *I_A_*. Juveniles mature into adults at rate *g* > 0 and adults reproduce at a maximum rate *a* > 0 subject to density-dependent competition given by *q* > 0 (juveniles do not reproduce). Juvenile and adult hosts die naturally at rates *b_J_* and *b_A_*. Disease transmission is assumed to be density-dependent, with stage-dependent transmission rates, *β_i_*(*r_i_*) = *β*_0_(1 – *r_i_*), where *β*_0_ > 0 is the baseline transmission rate and *r_i_* is host resistance at life-stage *i* (hence a host’s level of resistance determines the rate at which it becomes infected). Hosts are fully susceptible to infection when *r_i_* = 0 and fully resistant when *r_i_* = 1. The force of infection (rate at which susceptible hosts become infected) experienced at life-stage *i* is *λ_i_*(*r_i_*) = *β_i_*(*r_i_*)(*I_J_* + *I_A_*). We consider two types of virulence; infected hosts may either experience sterility virulence equal to 1 – *f*, where 0 ≤ *f* ≤ 1 is the reduction in fecundity when infected, or mortality virulence given by *α* > 0, the disease-associated mortality rate. We seek to compare the effects of mortality and sterility virulence and so we only allow the pathogen to exhibit one type of virulence at a time. We also assume that there is no recovery from infection, so that we can explore the effects of a lifelong reduction in fecundity on the evolution of juvenile resistance.

**Fig. 1:**
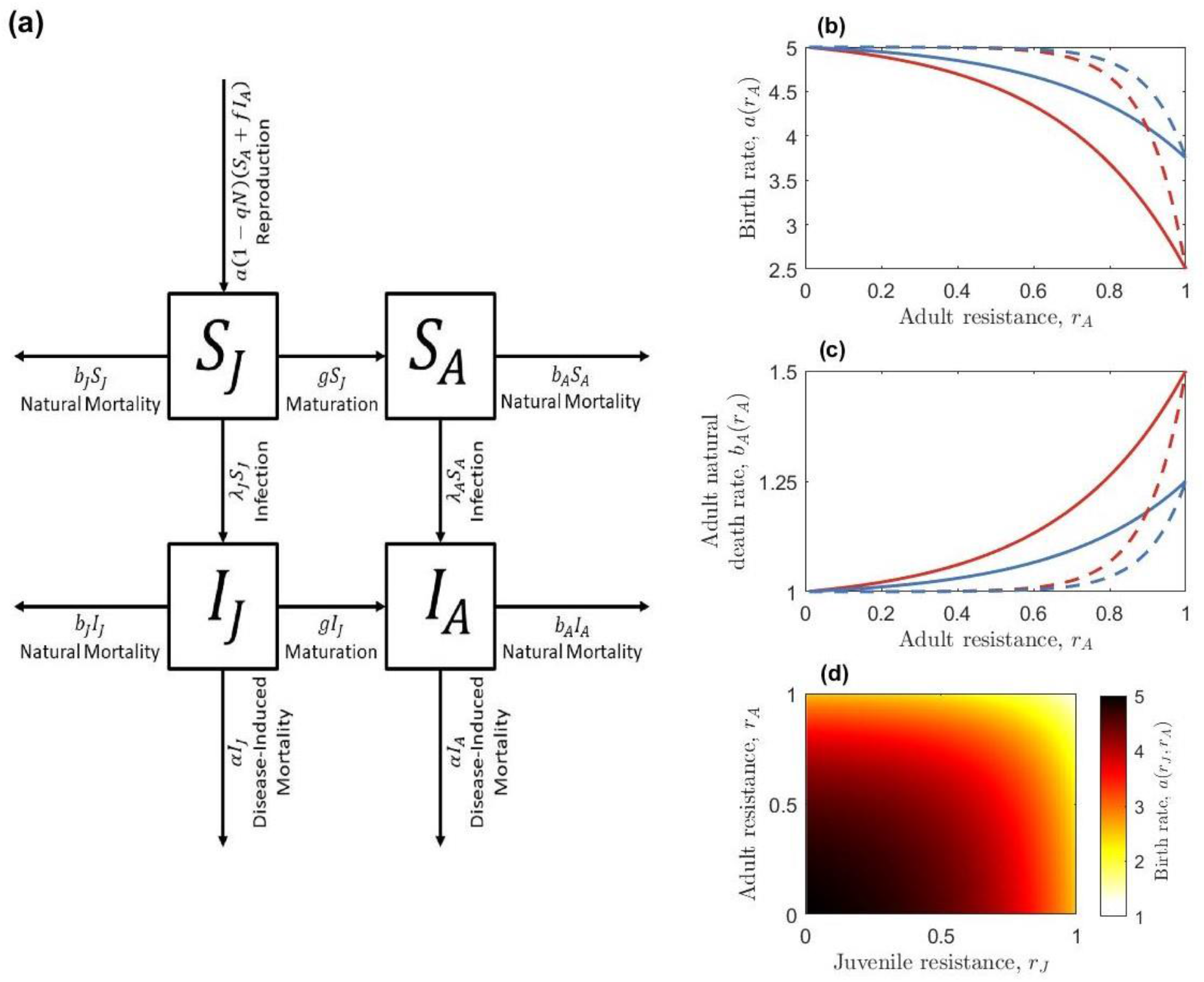
(a) Model schematic for the ecological model. (b-d) Examples of trade-off functions. Trade-offs are shown between: (b) adult resistance and birth rate (with *a*_0_ = 5), (c) adult resistance and adult mortality (with *b*_0_ = 1) and (d) both juvenile and adult resistance and the birth rate (with *a*_0_ = 5). Trade-offs between juvenile resistance and the maturation or birth rate take the same form as (b) and the trade-off between juvenile resistance and juvenile mortality takes the same form as (c). Trade-off strength is controlled by the parameter 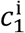; a relatively strong trade-off (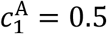, red curves) results in a much larger reduction in the birth rate for a given level of adult resistance than a relatively weak trade-off does (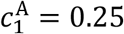, blue curves). Trade-off curvature is controlled by the parameter 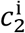; a relatively high curvature (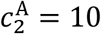, dashed lines) means that there is initially a low cost of increasing resistance but the cost eventually increases rapidly compared to a trade-off with lower curvature (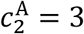, solid lines). Figure (d) is shown only in the strong, low curvature case.

**Table 1.** Model parameters and variables.

| Parameter/<br>variable | Description | Default value or<br>range |
| --- | --- | --- |
| $a$ | Reproduction rate of adult hosts | 5 |
| $b_J, b_A$ | Natural mortality rate of juvenile/adult hosts | 1 |
| $c_1^J, c_1^A$ | Strength of juvenile/adult trade-offs | 0.5 |
| $c_2^J, c_2^A$ | Curvature of juvenile/adult trade-offs | 3 |
| $1 - f$ | Sterility virulence | $0 \leq f \leq 1$ |
| $g$ | Host maturation rate | 1 |
| $I_J, I_A$ | Density of infected juveniles/adults | n/a |
| $N$ | Host population density | n/a |
| $q$ | Strength of host density-dependence | 1 |
| $r_J, r_A$ | Juvenile/adult resistance | $0 \leq r_J, r_A \leq 1$ |
| $S_J, S_A$ | Density of susceptible juveniles/adults | n/a |
| $t$ | Time, measured in arbitrary units | n/a |
| $\alpha$ | Mortality virulence | $0 \leq \alpha$ |
| $\beta_0$ | Baseline transmission rate | $0 \leq \beta_0$ |
| $\lambda_J, \lambda_A$ | Force of infection on juveniles/adults | n/a |

In a monomorphic population, the population dynamics are described by the following set of ordinary differential equations:

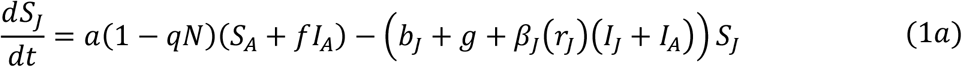

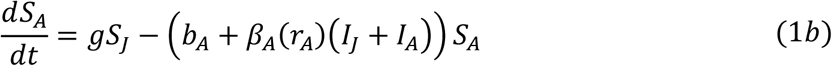

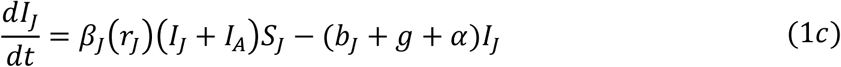

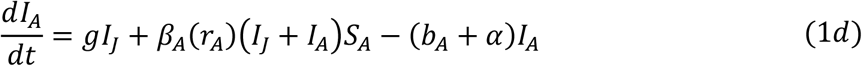

The disease-free equilibrium is given by:

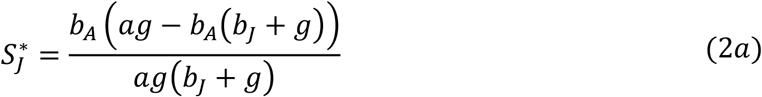

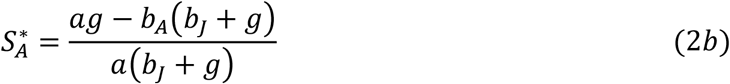

and is stable provided *ag* > *b_A_*(*b_J_* + *g*) and

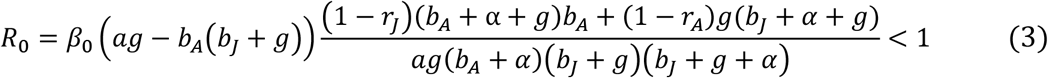

where *R*_0_ is the basic reproductive ratio of the pathogen (see *Supplementary Materials* for derivation). If (*b_J_* + *g*)*b_A_* > *ag* then the host population itself is not viable (individuals are lost from the juvenile and adult stages faster than they are added). The disease can spread when *R*_0_ > 1, in which case there is a stable, endemic (non-trivial) equilibrium for the parameters used in our analysis (this can be shown numerically, but there is no analytic expression for the endemic equilibrium; see *Supplementary Materials*).

In the absence of trade-offs, both juvenile and adult resistance will evolve to their maximum possible values (*r_J_*, *r_A_* = 1). We therefore assume that resistance at each life-stage trades off with another life-history trait. We consider a variety of trade-offs, with juvenile resistance either trading off with the maturation rate (*g*), reproduction rate (*a*) or juvenile mortality rate (*b_J_*) and adult resistance with either the reproduction rate (*a*) or adult mortality rate (*b_A_*). Biologically, these trade-offs assume that resistance requires hosts to divert resources from growth (slower maturation), reproduction (fewer offspring) or survival-related traits (higher mortality). We assume that resistance at each life-history stage only trades off with one other life-history trait. Specifically, we define the following trade-offs (when present) for the maturation rate,

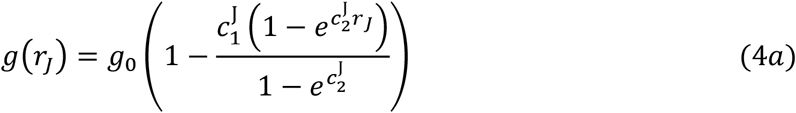

the reproduction rate, when it trades off with either juvenile (i = J) or adult (i = A) resistance

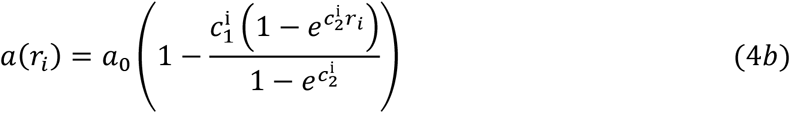

or with both juvenile and adult resistance,

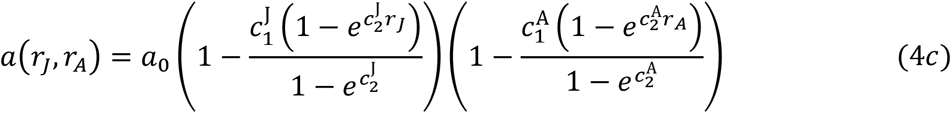

and the mortality rate

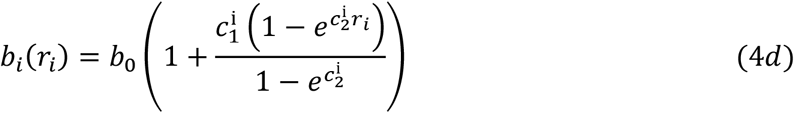

where *g*_0_, *a*_0_ and *b*_0_ are baseline maturation, reproduction and mortality rates (assuming equal baseline juvenile and adult mortality rates), 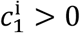 determines the maximum strength of the trade-off (i.e. the maximum proportional reduction or increase in the associated life-history trait) and 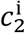 determines the curvature of the trade-off (larger absolute values correspond to greater deviations from linearity; Fig. 1b-d).

Intuitively, if the costs of resistance are sufficiently low at one life-stage relative to the other (e.g. 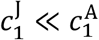) then resistance will always evolve to be higher at the life-stage with much lower costs. Hence one can easily choose trade-offs such that juvenile resistance is always greater than adult resistance, or vice versa. We therefore focus our analysis on how certain combinations of trade-offs promote higher juvenile or adult resistance, all else being equal, by keeping the proportional impact of all trade-offs the same 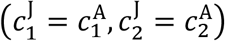, so that we can make fair comparisons across trade-offs. For example, if maximum juvenile resistance is associated with a 50% increase in juvenile mortality 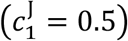, then we assume that maximum adult resistance is associated with either a 50% increase in adult mortality or a 50% decrease in reproduction 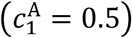. We only consider accelerating fitness costs 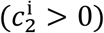, so that higher levels of resistance have diminishing returns, leading to evolutionarily stable strategies (decelerating fitness costs typically generate evolutionary repellers, but we restrict our attention to evolutionary attractors). We also fix the strength and curvature of the trade-offs such that 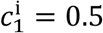 and 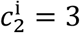, as our preliminary analysis revealed that variation in these parameters does not appear to cause qualitative changes to our key results (e.g., see Fig. S7, S8, S12, S13). It is also possible to rescale the system of equations (1a) to (1d) so that we can set *q* = 1 and *b*_0_ = 1 without loss of generality (see *Supplementary Materials*).

### Evolutionary Invasion Analysis

We use evolutionary invasion analysis (adaptive dynamics) to determine the coevolutionary dynamics of juvenile and adult resistance [52,53]. Specifically, we assume that mutations are sufficiently rare that there is a separation of ecological and evolutionary timescales (the ecological dynamics of the resident population reach equilibrium before the next mutation occurs) and that the mutations have small phenotypic effects. The invasion dynamics of rare host mutants are given in the *Supplementary Materials*. Using the next generation method [54], we derive the following expressions for the invasion fitness in the juvenile trait

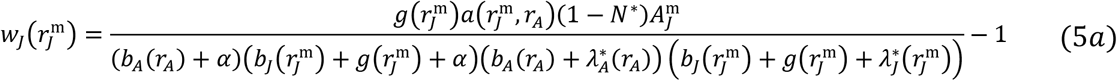

and in the adult trait

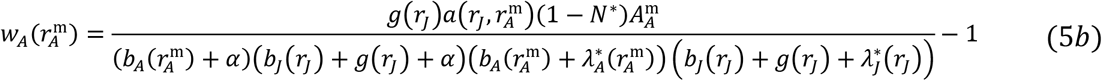

where asterisks denote the endemic equilibrium of the resident population. For notational convenience we set:

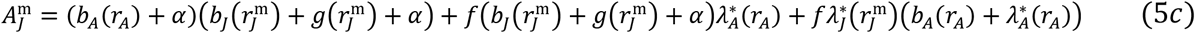

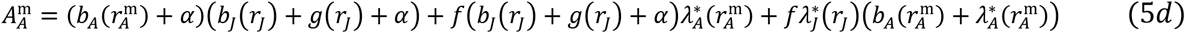

A mutant with juvenile resistance 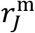 can invade a resident population (with resistance traits *r_J_* and *r_A_*) if and only if 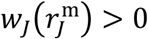, and similarly for a mutant with adult resistance 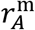. We assume equal mutation rates in juveniles and adults. There is no analytic expression for the endemic equilibrium of our model, so we cannot determine the singular strategies analytically. We therefore use numerical methods to calculate pairs of co-singular strategies (values of *r_J_* and *r_A_* that simultaneously maximise/minimise *w_J_* and *w_A_*) and to determine their evolutionary and strong convergence stability (see *Supplementary Materials*) [55,56]. Specifically, we calculate the fitness gradients and (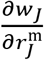 and 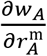 evaluated at 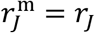 and 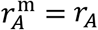 and solve simultaneously when both are equal to zero using numerical methods to give the co-singular strategies. We determine evolutionary stability by considering the signs of the second derivatives (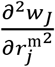 and 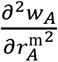 evaluated at the co-singular strategy). We determine strong convergence stability using other conditions on the second derivatives which tell us the signs of the real parts of the eigenvalues of the Jacobian matrix of the system (see *Supplementary Materials* for more details on the stability conditions). Evolutionary invasion analysis relies on the assumptions that mutations are rare and have small phenotypic effects. Also, strong convergence stability only guarantees that the co-singular strategy is an attractor of the evolutionary dynamics if the mutations have sufficiently small effects. We relax these assumptions by using evolutionary simulations to verify our results (see *Supplementary Materials* for a description of the simulations and for the source code).

## Results

### Sterility virulence

First, we consider the case where infection causes a reduction in the fecundity of the host (*f* < 1) but has no effect on host mortality (*α* = 0). From an ecological perspective, sterility virulence lowers reproduction and hence causes a reduction in the total population density. Such a reduction decreases the density of infected individuals and hence reduces the force of infection, reducing the density of infected individuals further than the density of susceptible individuals. This can be seen in Fig. 2 where the infected density initially falls in line with the total density, leaving the susceptible density relatively unchanged. Once sterility virulence rises sufficiently high, the host evolves resistance and the infected proportion falls further still.

**Fig. 2:**
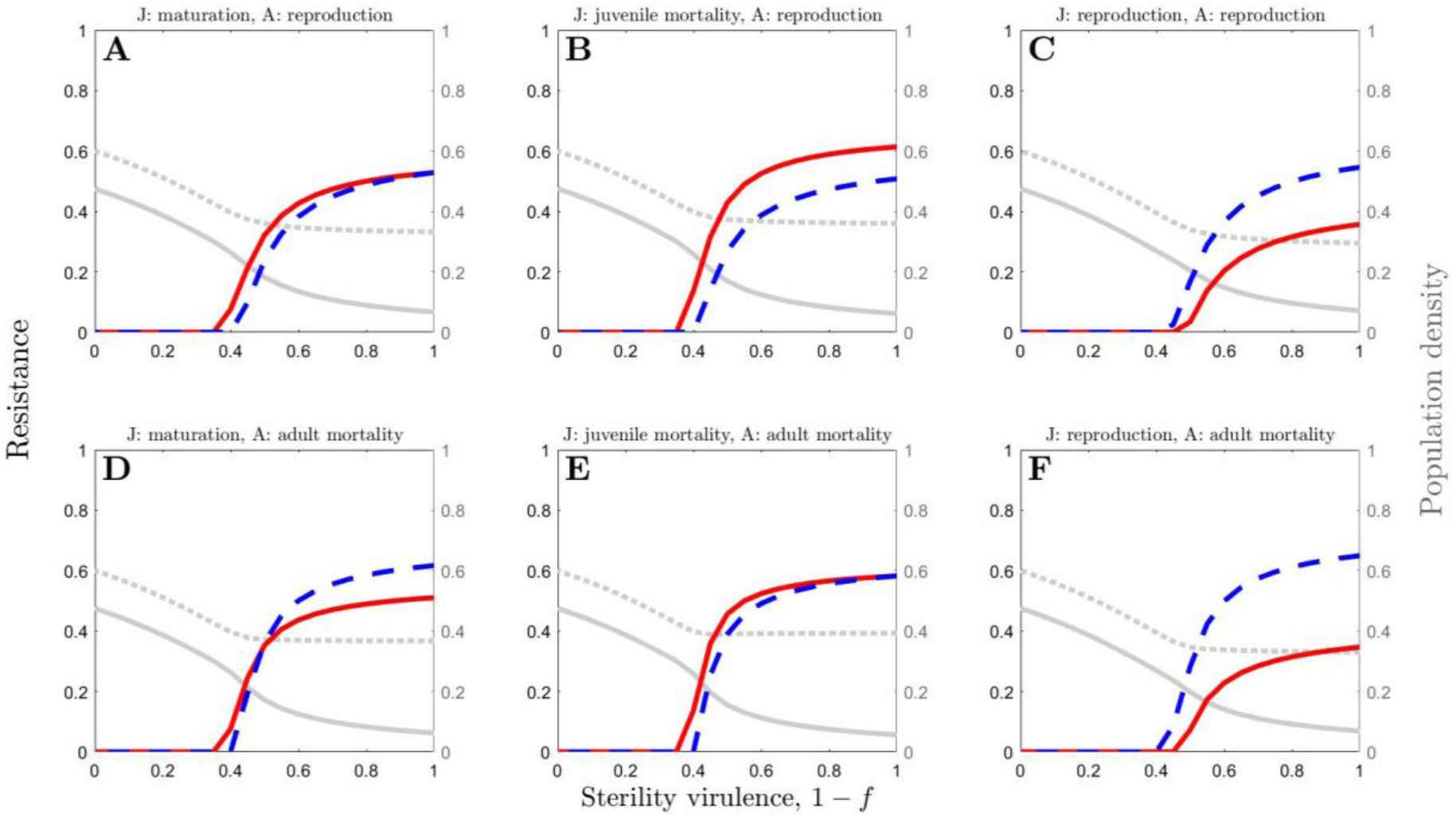
The effects of varying sterility virulence, 1 – *f*, on juvenile resistance (solid red) and adult resistance (dashed blue), for six different combinations of trade-offs: (A)-(C) adult resistance with reproduction, (D)-(F) adult resistance with adult mortality, (A) & (D) juvenile resistance with maturation, (B) & (D) juvenile resistance with juvenile mortality and (C) & (F) juvenile resistance with reproduction. The dotted grey line shows total population density and the solid grey line shows the density of infected hosts (both are non-dimensionalised). Parameter values are as in Table 1 with *β*_0_ = 8 and *α* = 0.

Unsurprisingly, neither adult nor juvenile resistance evolve for sufficiently low levels of sterility virulence but resistance at both life-stages may evolve when sterility virulence is sufficiently high (Fig. 2). Typically, juvenile and adult resistance both evolve towards a continuously stable strategy, although bistability is also possible for more extreme parameters (e.g., high transmissibility as shown in Fig. 3B and Fig. 4). We focus here on continuously stable strategies. If both juvenile and adult resistance are initially low then the density of infected individuals is likely to be relatively high and hence there may be selection for resistance at both life-stages. As both resistance traits increase, the number of infected individuals (and hence the risk of infection) falls, acting as a negative feedback on selection until both juvenile and adult resistance reach stable values (Fig. 3A). Similarly, if juvenile resistance is initially high and adult resistance is initially low, then small changes in adult resistance are far less costly than similar changes in juvenile resistance (because trade-offs are accelerating). Therefore, selection acts to decrease juvenile resistance (which results in a large reduction in costs) and increase adult resistance (which only leads to a small increase in costs).

**Fig. 3:**
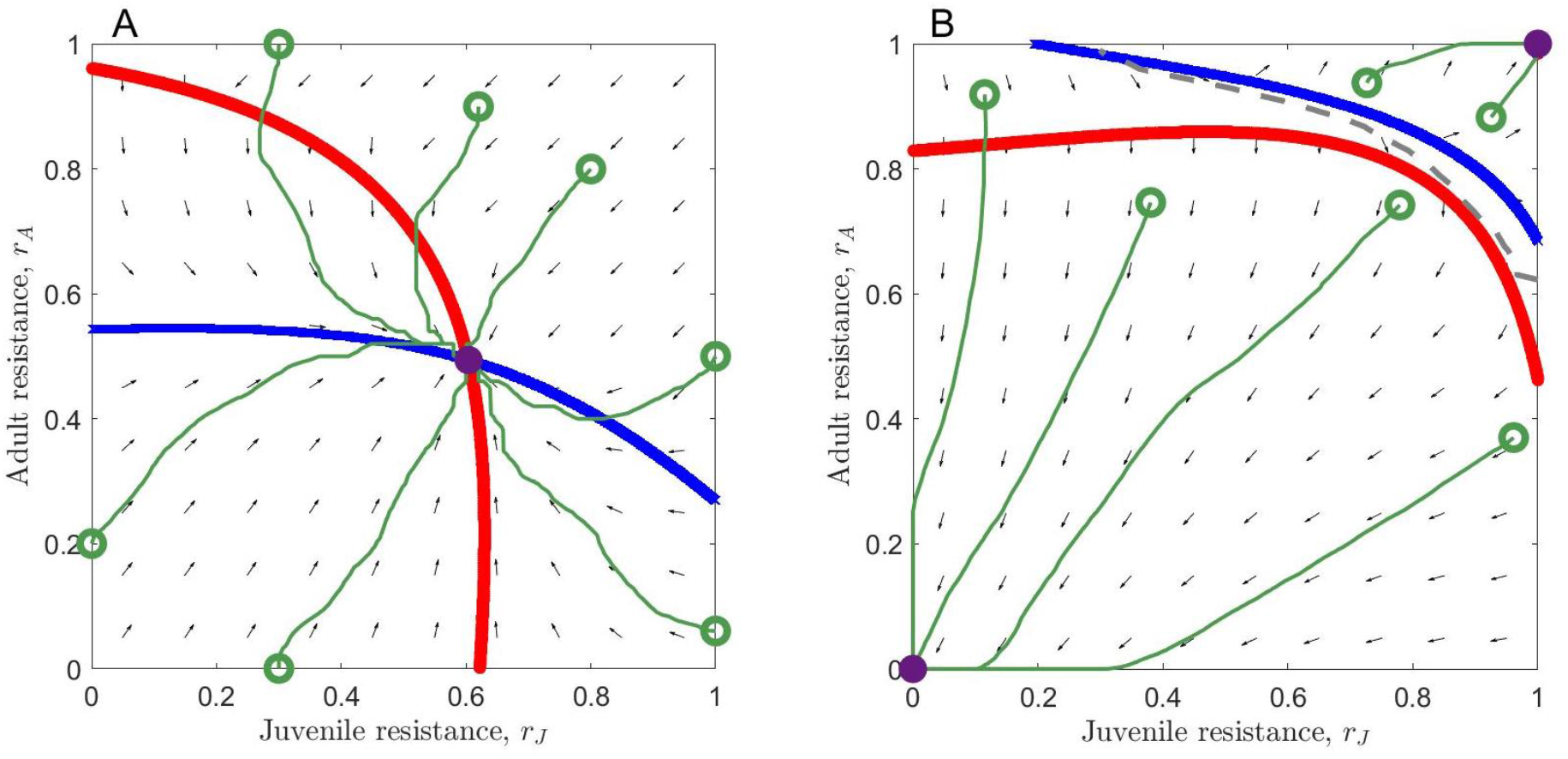
Phase planes showing (A) a continuously stable strategy and (B) bistability, with the juvenile nullcline in red and the adult nullcline in blue. In (A), the host population will always evolve towards the CSS (purple circle), no matter what the starting values of the juvenile and adult resistance traits. In (B), the host population will evolve towards one of the attractors (purple circles), depending on the starting values of the juvenile and adult resistance traits (basins of attraction are separated by the dashed line). Example trajectories are shown in green. In (A), juvenile resistance trades off with juvenile mortality, adult resistance trades off with reproduction and parameter values are as in Table 1 with *β*_0_ = 8, *α* = 0 and *f* = 0.1. In (B), juvenile resistance trades off with juvenile mortality, adult resistance trades off with adult mortality and parameter values are as in Table 1 with *β*_0_ = 1000 (high transmissibility), *α* = 0 and *f* = 0.5.

**Fig. 4:**
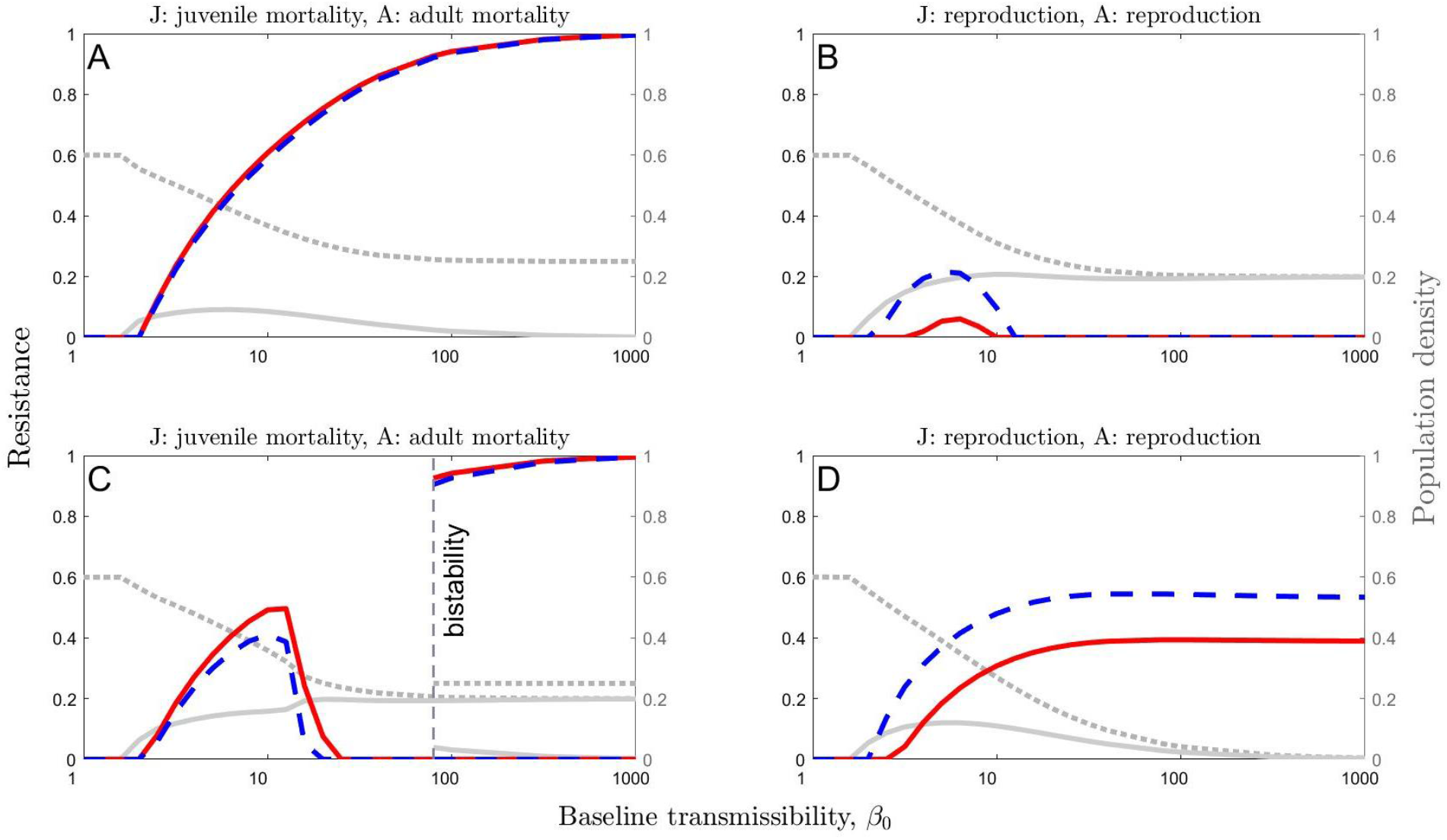
The effect of varying baseline transmissibility, *β*_0_, on juvenile resistance (solid red) and adult resistance (dashed blue), in the cases where juvenile resistance trades off with juvenile mortality and adult resistance trades off with adult mortality (A and C) and where both juvenile and adult resistance trade off with reproduction (B and D). The dotted grey line shows total population density and the solid grey line shows the density of infected hosts (both are non-dimensionalised). In the bistability region in panel C, the higher total population density and the lower infected density correspond to the higher levels of resistance. Parameter values are as in Table 1, with *α* = 0 and *f* = 0.5 (B and C) or *f* = 0.3 (A and D).

The stable levels of juvenile and adult resistance will clearly depend on the nature of the trade-offs involved, as equal levels of resistance will generally not incur the same cost to the host. However, regardless of which life-history traits trade-off with resistance and at which life-stage resistance acts, the general shape of the resistance curve in response to variation in sterility virulence is consistent. Specifically, at moderate levels of sterility virulence there is a sharp increase in resistance but this plateaus when sterility virulence is high. This suggests that when sterility virulence is at moderate levels, a relatively small increase in virulence can lead to a marked increase in selection for resistance at both juvenile and adult stages, regardless of the underlying trade-offs.

All else being equal (i.e. trade-offs have the same proportional effect on life-history traits for a given level of resistance), juvenile and adult resistance are typically similar if juvenile resistance trades off with maturation (Fig. 2A, D) or if resistance is associated with an increase in mortality (Fig. 2E). If, however, juvenile resistance is associated with higher juvenile mortality and adult resistance is associated with lower reproduction, our model predicts that juvenile resistance is consistently higher than adult resistance (Fig. 2B). Conversely, if juvenile resistance trades off with adult reproduction, then adult resistance is consistently higher than juvenile resistance regardless of whether adult resistance trades off with reproduction (Fig. 2C) or mortality (Fig. 2F), and we also see lower levels of juvenile resistance than we do when other trade-offs are present (Fig. 2, S1). Since there is no recovery in our model, becoming infected as a juvenile leads to a permanent reduction in fecundity, yet our model suggests that risking infection as a juvenile is generally a better strategy than investing in resistance if this incurs a reproduction cost. In the *Supplementary Materials*, we show how these results can be interpreted using the trade-off terms in the fitness gradients. For example, in the case of both resistance traits trading off with reproduction, we show that selection for lower resistance at the juvenile stage occurs due to less time spent, on average, as a susceptible juvenile than as a susceptible adult. Hence the benefits of juvenile resistance are lower than for adult resistance.

These results are qualitatively consistent for variation in the baseline reproduction (*a*_0_) and maturation (*g*_0_) rates and trade-off parameters (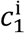 and 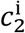)(Fig. S9-S13), with adult resistance exceeding juvenile resistance most markedly when maturation is fast and when juvenile resistance trades off with reproduction (Fig. S10C and S10F).

Similarly, variation in baseline transmissibility (*β*_0_) affects the risk of infection for adults and juveniles equally and so has a similar effect on resistance evolution at both life-stages (Fig. 4). When *β*_0_ is very low, the risk of infection is low and hence resistance does not evolve at either life-stage. As *β*_0_ increases, the disease becomes more common, leading to a higher infected density and a smaller total host population (due to sterility virulence). In evolutionary terms, this also causes both juvenile and adult resistance to rise (Fig. 4), with similar differences between trade-offs as described above (Fig. S2 and Fig. S3). For sufficiently high values of *β*_0_, the outcome depends on whether the host population remains viable (see *Supplementary Materials*), in which case resistance may tend towards either a high value if the pathogen is sufficiently virulent (Fig. 4A) or else a low value if disease prevalence approaches 100% with most individuals infected very shortly after birth (with selection against ineffective resistance, as observed in some empirical systems [57]; Fig. 4B).

Alternatively, for some parameter and trade-off combinations, the population may enter a region of bistability for extremely high values of *β*_0_ (Fig. 4C), where hosts either evolve to high or zero levels of resistance at both life-stages, depending on the initial levels of resistance in the population (Fig. 3B). If the resistance traits are initially low then the high pathogen transmissibility will produce a very high infected density. This means that hosts will become infected almost immediately after birth and so resistance (other than an extremely high level of resistance) will be ineffective. Therefore, selection acts against ineffective resistance. If, however, the resistance traits are initially very high then this will suppress the pathogen. Any reduction in resistance will be detrimental for the host as it will allow the infected density to rise significantly and so selection acts for increased resistance. This bistability suggests that, in principle, initially similar populations could experience very different evolutionary outcomes, although such high levels of transmissibility are unlikely to be biologically realistic. Finally, if the host population size tends towards zero as *β*_0_ increases, then resistance tends towards an intermediate level (e.g. Fig. 4D), although the level of resistance is inconsequential as the host population crashes.

### Mortality Virulence

We now consider the case where infection increases the mortality rate (*α* > 0) but has no effect on host fecundity (*f* = 1). From an ecological perspective, population density, particularly of infected individuals, falls as mortality virulence increases, but sufficiently high mortality virulence initiates the evolution of resistance. Juvenile and adult resistance follow the same qualitative patterns as mortality virulence varies. As in non-age-structured models, hosts do not evolve resistance when *α* is sufficiently low because the costs of infection are low, or when *α* is sufficiently high because this reduces the infectious period and hence lowers the density of infected individuals at any given time. Resistance therefore peaks at intermediate values of *α*, although both the extent of resistance and when it peaks may differ between life-stages (Fig. 5). Moreover, certain combinations of trade-offs consistently favour higher juvenile resistance and others higher adult resistance, all else being equal (Fig. 5). Specifically, juvenile resistance tends to be markedly lower than adult resistance when the former trades off with maturation or natural mortality rate (Fig. 5A-B, D-E) but the converse is true when juvenile resistance trades off with adult reproduction (Fig. 5C, F). We can see that juvenile resistance is significantly lower in the latter case (Fig. 5C, F) than in the former cases (Fig. 5A-B, D-E). These patterns are consistent as other model parameters are varied (Fig. S4-S8) and largely mirror those for sterility virulence (Fig. 2).

**Fig. 5:**
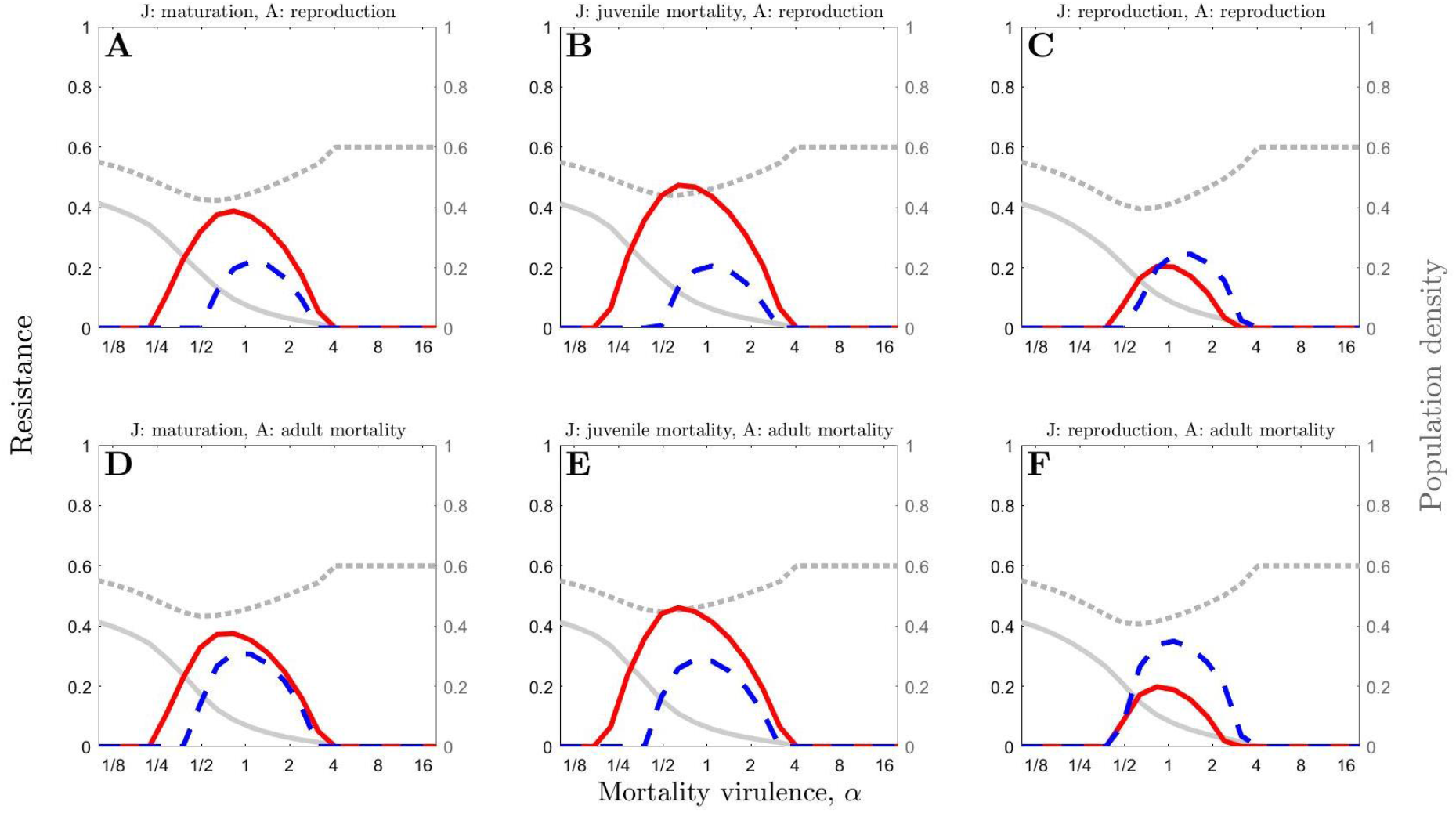
The effect of varying mortality virulence, *α*, on juvenile resistance (solid red) and adult resistance (dashed blue), for six different combinations of trade-offs: (A)-(C) adult resistance with reproduction, (D)-(F) adult resistance with adult mortality, (A) & (D) juvenile resistance with maturation, (B) & (D) juvenile resistance with juvenile mortality and (C) & (F) juvenile resistance with reproduction. The dotted grey line shows total population density and the solid grey line shows the density of infected hosts (both are non-dimensionalised). Parameter values are as in Table 1 with *β*_0_ = 8 and *f* = 1.

## Discussion

Significant differences in innate, infection-preventing resistance have been observed between juveniles and adults across many taxa and yet the evolutionary drivers of these differences are not well understood [51]. Here, we theoretically explored how trade-offs between age-specific resistance and various life-history traits combine to affect selection for resistance at different life-stages and investigated whether selection typically favours higher juvenile or adult resistance, all else being equal. Overall, our analysis suggests that trade-offs between juvenile resistance and adult reproduction are inherently more costly than other trade-offs, regardless of whether virulence affects mortality or fecundity. These particular trade-offs may lead to the evolution of relatively low resistance as a juvenile (compared with adult resistance and with juvenile resistance when other trade-offs are present), even when infection as a juvenile causes lifelong reductions in fecundity. The latter result may appear counter-intuitive at first, but if the lifelong reduction in fecundity due to infection and the risk of infection as a juvenile are both sufficiently low, then it is better for the host to risk infection as a juvenile rather than always to suffer from a reduced reproduction rate as an adult.

We fixed the strength and shape of the trade-offs in our model to be the same for all tradeoff functions so that we could make fair comparisons between different combinations of trade-offs. Hence, our key finding that adult resistance tends to be relatively higher when juvenile resistance trades off with reproduction suggests that this is because it is inherently costlier, compared to trade-offs with maturation or mortality, for hosts to evolve juvenile resistance if it results in decreased reproduction as an adult. This also suggests that costs of juvenile resistance associated with reproduction may have a disproportionately greater effect on host fitness than costs associated with maturation or mortality. Whether juvenile resistance is higher than adult resistance, or vice versa, in a particular host-pathogen system will also depend on the strength and shape of the trade-offs. For example, if a given level of adult resistance is proportionately much more costly than a given level of juvenile resistance, then we should expect juvenile resistance to be higher. However, we predict that when the strength and shape of the trade-offs are similar, adult resistance will tend to be higher than juvenile resistance if the latter trades off with reproduction. This result may also provide clues as to where trade-offs may exist if empirical observations reveal that juveniles are intrinsically less resistant than adults.

Our study examined the effect of trade-offs with different life-history traits: mortality, maturation and fecundity. In plants, where costs of resistance have been relatively well-studied, trade-offs between innate, infection-preventing resistance and fecundity are well supported [42–44,58–61]. In many crop plants, resistance is typically measured at the seedling (juvenile) stage whereas costs may be measured in growing or mature (adult) plants. For example, in oats, seedling resistance to infection by rust fungi has been linked to substantial (9%) reductions in yield [59]. In tobacco, resistance to infection by tobacco mosaic virus, measured at 4 weeks post planting, led to reduced growth [61]. In *Arabidopsis*, a resistance gene that affects the ability of a bacterial pathogen to invade at 3 weeks of age (when plants are in the young rosette stage), has been associated with up to 9% reductions in seed set [60]. There is also some evidence of costs associated with maturation rate. For example, Barlett *et al*. found a negative correlation between maturation rate and resistance to infection by a baculovirus at the third-instar larval stage in the moth *plodia interpunctella* [45]. Survival is less commonly investigated as a potential trade-off mechanism and there is currently little evidence for trade-offs between survival and innate resistance (although see [62] for a review of immunopathology). Our study shows that when costs are paid through reductions in fecundity, adult resistance is favoured over juvenile resistance in most cases.

It is critical to note that whilst trade-offs have been documented for both juvenile and adult resistance, we can find no study that directly quantifies the magnitude of these costs within a single host. This is largely because resistance phenotyping is typically done at a single age, or in the case of crop studies, seedling and adult resistance are measured in completely different settings with different inoculum sources and so are difficult to compare [63–65]. One study by Biere & Antonovics found a negative correlation between flower production and resistance of adult *Silene latifolia* plants to anther-smut infection in a field setting, but no apparent correlation between flower production and family-level resistance measured in the lab at the seedling stage [42]. It is, however, reasonable to expect (from a resource allocation perspective) that diversion of resources to resistance during development could negatively impact on adult fecundity, for instance by restricting growth (body size or secondary sex traits) which could make individuals less competitive for mates or less able to support a larger number of offspring. Our results demonstrate that quantifying the magnitude and form of such trade-offs at juvenile and adult stages is critically important for determining the evolutionary outcomes of age-specific resistance. We tentatively predict that, in systems where juveniles are less resistant to infection than adults, trade-offs between juvenile resistance and reproduction may be more likely than trade-offs between other life-history traits.

This prediction could be tested using a host species which is naturally polymorphic in resistance to a particular pathogen. Having bred separate families of hosts, the juvenile and adult resistance of each family could be estimated by exposing hosts of different ages to the pathogen and calculating the proportion of each age-group within each family which becomes infected. Other individuals from each family could be used to measure possible trade-off traits at different life-stages (for instance growth or reproduction). A negative correlation between resistance at any life-stage and any other beneficial trait would suggest a trade-off.

Our results are broadly consistent as our model parameters are varied, although when the pathogen is highly transmissible it is possible for the host to experience bistability, with selection either favouring high juvenile and adult resistance or no resistance across the life span, depending on the initial conditions. This suggests that founder effects, or drift reinforced by selection, could drive initially similar populations to contrasting evolutionary outcomes. However, we found no evidence of bistability causing levels of resistance to diverge substantially at different life-stages (i.e. high juvenile resistance and no adult resistance, or vice versa). Bistability is therefore not likely to be the cause of contrasting levels of resistance in juveniles and adults.

Previous theory has almost entirely focused on the evolution of resistance in populations without age-structure [46–50]. Our model was an extension of the one explored by Ashby & Bruns, which considered the evolution of juvenile susceptibility (the inverse of resistance) subject to trade-offs with reproduction or maturation [51]. However, Ashby & Bruns assumed that hosts were always more resistant as adults than as juveniles [51], whereas here we have relaxed these assumptions to consider how juvenile and adult resistance evolve simultaneously subject to a wider range of trade-offs.

We made several simplifying assumptions in the process of modelling this evolutionary process. Firstly, we assumed that juvenile and adult resistance evolve independently, which is reasonable if different mechanisms are responsible for resistance at different life-stages [66], but instead juvenile and adult resistance may be correlated if the mechanism is the same. Secondly, we assumed that each resistance trait only incurred one type of cost rather than trading off with multiple life-history traits, which is reasonable from a general modelling perspective, but may not hold true in certain systems where, for example, juvenile resistance may trade-off against multiple life-history traits such as maturation, reproduction and mortality. Thirdly, we assumed that disease effects on juveniles and adults were identical, but the severity of disease may differ depending on the age of the host. For example, age is a strong predictor of the risk of mortality from COVID-19 in humans [67]. Including age-related disease effects in our model would have greatly complicated our analysis, but this should be considered in future theoretical work. Similarly, we assumed that juveniles and adults mixed randomly, but the effects of biased (assortative) transmission between individuals at the same life-stage should also be considered in future work.

Fourthly, we assumed that there was no recovery from infection, as our model was loosely inspired by the sterilising anther-smut pathogen (*Microbotryum*) in carnations (*Caryophyllaceae*), which rarely recover from infection but exhibit substantial variation in resistance between seedling and mature plants [68]. Preliminary analysis revealed that recovery from infection does not change our key results, but by assuming that there was no recovery we were readily able to explore the effects of lifelong reductions in fecundity arising from infection as a juvenile. Finally, we assumed that the pathogen was monomorphic and evolutionarily static. Clearly, in a real-world scenario the pathogen would be expected to evolve in response to changes in the host and so future models should consider the effects of host-pathogen coevolution in age-structured populations. This could include the evolution of either parasite infectivity or virulence, which would also extend previous theoretical work on the evolution of stage-specific virulence [41]. Host-pathogen coevolution with age-specific resistance has yet to be explored theoretically [69].

In our model, we focused on the evolution of innate, infection-preventing resistance, as opposed to other forms of host defence such as tolerance. Both forms of defence against pathogens are common in nature, with resistance and tolerance strategies operating concurrently in many cases. However, age-structured tolerance is not well-understood and would therefore be difficult to model. For instance, how would the host’s level of tolerance change as it aged from a juvenile to an adult whilst infected? Combining the two types of defence might also complicate matters if resistance and tolerance had significant effects on one another. Future work should consider how tolerance may evolve across the lifespan of the host.

Overall, our model shows that trade-offs between juvenile resistance and reproduction during adulthood are intrinsically more costly than trade-offs between other traits, even when infection leads to permanent reductions in fecundity. Such trade-offs could therefore explain why adults are sometimes more resistant to disease than juveniles.

## Acknowledgements

We thank Nick Priest for helpful discussions about the manuscript.

## Data Accessibility Statement

Source code is available in the *Supplementary Materials* and at https://github.com/ecoevotheory/Buckingham_and_Ashby_2022.

## Funding

Ben Ashby is supported by the Natural Environment Research Council (grant nos. NE/N014979/1 and NE/V003909/1). This research was generously supported by a Milner Scholarship PhD grant to Lydia Buckingham from The Evolution Education Trust.

## Notes

https://github.com/ecoevotheory/Buckingham_and_Ashby_2022

### Competing Interest Statement

The authors have declared no competing interest.

### Summary of Updates

This version of the manuscript has been revised in response to comments from reviewers. Changes to the main text are minor. An additional section has been added to the Supplementary Materials.

