## Supplementary Materials for "The evolution of age-specific resistance to infectious disease"

### The evolution of age-specific resistance to infectious disease: Supplementary Materials

#### Non-Dimensionalisation

We scale the system of equations (1a) to (1d) in the main text as follows:

$$S_J, S_A, I_J, I_A, N \sim \frac{1}{q} \qquad (S1a)$$

$$t \sim \frac{1}{b_0} \qquad (S1b)$$

$$a, g, \alpha \sim b_0 \qquad (S1c)$$

$$\beta_0 \sim q b_0 \qquad (S1d)$$

If we then divide the whole system by $\frac{b_0}{q}$, it returns to our original system of equations (1a) to (1d) but with $q = 1$ and $b_0 = 1$.

This allows us to set $q = 1$ and $b_0 = 1$ without loss of generality.

#### Derivation of $R_0$

The dynamics of a small number of infected individuals in a population close to its disease-free equilibrium are given by:

$$\frac{\mathrm{d}I_J}{\mathrm{d}t} = \lambda_J S_J^* - (b_J + g + \alpha) I_J \qquad (S2a)$$

$$\frac{\mathrm{d}I_A}{\mathrm{d}t} = g I_J + \lambda_A S_A^* - (b_A + \alpha) I_A \qquad (S2b)$$

The next-generation matrix [1] of this system may be written as:

$$N_G = \frac{\beta_0}{(b_A + \alpha)(b_J + g + \alpha)} \begin{pmatrix} (1 - r_J)(hb_A + h\alpha + g) S_J^* & (1 - r_J)(b_J + g + \alpha) S_J^* \\ (1 - r_A)(hb_A + h\alpha + g) S_A^* & (1 - r_J)(b_J + g + \alpha) S_A^* \end{pmatrix} \qquad (S3)$$

The largest eigenvalue of this matrix gives the basic reproductive ratio:

$$R_0 = \beta_0 \left(ag - b_A(b_J + g)\right) \frac{(1 - r_J)(hb_A + h\alpha + g)b_A + (1 - r_A)g(b_J + \alpha + g)}{ag(b_A + \alpha)(b_J + g)(b_J + g + \alpha)} \qquad (S4)$$

**POPULATION VIABILITY**

First consider a host population with no resistance ($r_J = r_A = 0$). In the absence of disease, our system becomes:

$$\frac{\mathrm{d}S_J}{\mathrm{d}t} = a_0(1 - S_J - S_A)S_A - (g_0 + b_0)S_J \qquad (S5a)$$

$$\frac{\mathrm{d}S_A}{\mathrm{d}t} = g_0 S_J - b_0 S_A \qquad (S5b)$$

which has a single, non-trivial (stable) equilibrium at:

$$S_A^* = \frac{a_0 g_0 - b_0(g_0 + b_0)}{a_0(g_0 + b_0)} \qquad (S6a)$$

$$S_J^* = \frac{b_0}{g_0} S_A^* \qquad (S6b)$$

Hence, a disease-free population with no resistance is viable if $a_0 g_0 > b_0(g_0 + b_0)$.

Now consider the case where individuals become infected as soon as they are born, which can be approximated by:

$$\frac{\mathrm{d}I_J}{\mathrm{d}t} = a_0(1 - I_J - I_A)fI_A - (g + b_0 + \alpha)I_J \qquad (S7a)$$

$$\frac{\mathrm{d}I_A}{\mathrm{d}t} = g_0 I_J - (b_0 + \alpha)I_A \qquad (S7b)$$

which has a single non-trivial (stable) equilibrium at:

$$I_A^* = \frac{a_0 f g_0 - (b_0 + \alpha)(g_0 + b_0 + \alpha)}{a_0 f (g_0 + b_0 + \alpha)} \qquad (S8a)$$

$$I_J^* = \frac{b_0 + \alpha}{g_0} I_A^* \qquad (S8b)$$

Therefore, an entirely infected population is viable if $a_0 f g_0 > (b_0 + \alpha)(g_0 + b_0 + \alpha)$.

Suppose instead that the host population is completely resistant $(r_J = r_A = 1)$. In this case, the disease cannot spread and so our system reduces to:

$$\frac{\mathrm{d}S_J}{\mathrm{d}t} = a(1 - S_J - S_A)S_A - (g + b_J)S_J \qquad (S9a)$$

$$\frac{\mathrm{d}S_A}{\mathrm{d}t} = gS_J - b_A S_A \qquad (S9b)$$

where $a$, $g$, $b_J$ and $b_A$ are evaluated at $r_J = r_A = 1$, where appropriate. A stable, non-trivial equilibrium exists at:

$$S_A^* = \frac{ag - b_A(g + b_J)}{a(g + b_J)} \qquad (S10a)$$

$$S_J^* = \frac{b_A}{g} S_A^* \qquad (S10b)$$

Therefore, an entirely resistant population is viable if $ag > b_A(g + b_J)$ at $r_J = r_A = 1$.

##### STABILITY OF DISEASE-FREE ECOLOGICAL EQUILIBRIUM

In the disease-free case, our system of equations reduces to:

$$\frac{\mathrm{d}S_J}{\mathrm{d}t} = a(1 - S_J - S_A)S_A - (g + b_J)S_J \qquad (S11a)$$

$$\frac{\mathrm{d}S_A}{\mathrm{d}t} = gS_J - b_A S_A \qquad (S11b)$$

The disease-free equilibrium of this system is given by:

$$S_J^* = \frac{b_A\left(ag - b_A(b_J + g)\right)}{ag(b_J + g)} \qquad (S12a)$$

$$S_A^* = \frac{ag - b_A(b_J + g)}{a(b_J + g)} \qquad (S12b)$$

We can linearise the system about this equilibrium by setting $S_J = S_J^* + \hat{S}_J$ and $S_A = S_A^* + \hat{S}_A$ where $\hat{S}_J$ and $\hat{S}_A$ are small deviations from the equilibrium. Substituting these expressions into the system above and neglecting quadratic and higher order terms in $\hat{S}_J$ and $\hat{S}_A$, we get:

$$\frac{\mathrm{d}\widehat{S_J}}{\mathrm{d}t} = a(1 - S_J^* - S_A^*)\hat{S}_A + a(-\hat{S}_J - \hat{S}_A)S_A^* - (g + b_J)\hat{S}_J \qquad (S13a)$$

$$\frac{\mathrm{d}\hat{S}_A}{\mathrm{d}t} = g\hat{S}_J - b_A\hat{S}_A \qquad (S13b)$$

This system can be written in matrix form as:

$$\frac{\mathrm{d}}{\mathrm{d}t}\begin{pmatrix}\hat{S}_J \\ \hat{S}_A\end{pmatrix} = M\begin{pmatrix}\hat{S}_J \\ \hat{S}_A\end{pmatrix} \qquad (S14a)$$

where:

$$M = \begin{pmatrix} -aS_A^* - b_J - g & a(1 - S_J^* - S_A^*) - aS_A^* \\ g & -b_A \end{pmatrix} \qquad (S14b)$$

The equilibrium point is linearly stable if $\mathrm{tr}(M) < 0$ and $\det(M) > 0$.

We can clearly see that $\mathrm{tr}(M) = -aS_A^* - b_J - g - b_A < 0$.

Substituting expressions (S12) into $\det(M) = b_A(aS_A^* + b_J + g) + g(2aS_A^* + aS_J^* - a)$ gives:

$$\det(M) = (2b_A - b_J + g)aS_A^* \tag{S15}$$

which is positive because we only ever consider $c_1^i < 1$ and so we always have $b_J < 2b_A$.

Therefore, the disease-free equilibrium is linearly stable.

#### STABILITY OF ENDEMIC ECOLOGICAL EQUILIBRIUM

We can linearise the full system about its endemic equilibrium ($S_J^*$, $S_A^*$, $I_J^*$ and $I_A^*$) and into the form:

$$\frac{\mathrm{d}}{\mathrm{d}t}\begin{pmatrix}\hat{S}_J\\ \hat{S}_A\\ \hat{I}_J\\ \hat{I}_A\end{pmatrix} = M\begin{pmatrix}\hat{S}_J\\ \hat{S}_A\\ \hat{I}_J\\ \hat{I}_A\end{pmatrix} \tag{S16a}$$

where:

$$M = \begin{pmatrix} -a(S_A^* + fI_A^*) - (b_J + g + \lambda_J^*) & a(1 - N^*) - a(S_A^* + fI_A^*) & -a(S_A^* + fI_A^*) - \beta_J S_J^* & a(1 - N^*) - a(S_A^* + fI_A^*) - \beta_J S_J^* \\ g & -(b_A + \lambda_A^*) & -\beta_A S_A^* & -\beta_A S_A^* \\ \lambda_J^* & 0 & \beta_J S_J^* - (b_J + \alpha + g) & \beta_J S_J^* \\ 0 & \lambda_A^* & g + \beta_A S_A^* & \beta_A S_A^* - (b_A + \alpha) \end{pmatrix} \tag{S16b}$$

The endemic equilibrium is linearly stable if the real part of each of the eigenvalues of $M$ is negative. We cannot calculate the eigenvalues of this matrix analytically and so we do this numerically for different sets of parameter values (see source code). We find that the endemic equilibrium is unique and linearly stable whenever it exists, across a wide range of parameter values.

#### EVOLUTIONARY INVASION ANALYSIS

The invasion dynamics of a rare host mutant with juvenile resistance $r_J^{\mathrm{m}}$ in an established resident population (at the endemic equilibrium, denoted by asterisks) in the case where juvenile resistance trades off with the maturation rate, $g(r_J^{\mathrm{m}})$, and adult resistance trades off with the birth rate, $a(r_A)$, are given by:

$$\frac{\mathrm{d}S_J^{\mathrm{m}}}{\mathrm{d}t} = a(r_A)(1 - N^*)(S_A^{\mathrm{m}} + fI_A^{\mathrm{m}}) - \left(b_J + g\left(r_J^{\mathrm{m}}\right) + \lambda_J^*\left(r_J^{\mathrm{m}}\right)\right)S_J^{\mathrm{m}} \qquad (S17a)$$

$$\frac{\mathrm{d}S_A^{\mathrm{m}}}{\mathrm{d}t} = g\left(r_J^{\mathrm{m}}\right)S_J^{\mathrm{m}} - \left(b_A + \lambda_A^*(r_A)\right)S_A^{\mathrm{m}} \qquad (S17b)$$

$$\frac{\mathrm{d}I_J^{\mathrm{m}}}{\mathrm{d}t} = \lambda_J^*\left(r_J^{\mathrm{m}}\right)S_J^{\mathrm{m}} - \left(b_J + g\left(r_J^{\mathrm{m}}\right) + \alpha\right)I_J^{\mathrm{m}} \qquad (S17c)$$

$$\frac{\mathrm{d}I_A^{\mathrm{m}}}{\mathrm{d}t} = g\left(r_J^{\mathrm{m}}\right)I_J^{\mathrm{m}} + \lambda_A^*(r_A)S_A^{\mathrm{m}} - (b_A + \alpha)I_A^{\mathrm{m}} \qquad (S17d)$$

Similar equations can be obtained for other combinations of trade-offs. Likewise, the dynamics of a rare mutant with adult resistance $r_A^{\mathrm{m}}$ are given by:

$$\frac{\mathrm{d}S_J^{\mathrm{m}}}{\mathrm{d}t} = a(r_A^{\mathrm{m}})(1 - N^*)(S_A^{\mathrm{m}} + fI_A^{\mathrm{m}}) - \left(b_J + g\left(r_J\right) + \lambda_J^*\left(r_J\right)\right)S_J^{\mathrm{m}} \qquad (S18a)$$

$$\frac{\mathrm{d}S_A^{\mathrm{m}}}{\mathrm{d}t} = g\left(r_J\right)S_J^{\mathrm{m}} - \left(b_A + \lambda_A^*(r_A^{\mathrm{m}})\right)S_A^{\mathrm{m}} \qquad (S18b)$$

$$\frac{\mathrm{d}I_J^{\mathrm{m}}}{\mathrm{d}t} = \lambda_J^*\left(r_J\right)S_J^{\mathrm{m}} - \left(b_J + g\left(r_J\right) + \alpha\right)I_J^{\mathrm{m}} \qquad (S18c)$$

$$\frac{\mathrm{d}I_A^{\mathrm{m}}}{\mathrm{d}t} = g\left(r_J\right)I_J^{\mathrm{m}} + \lambda_A^*(r_A^{\mathrm{m}})S_A^{\mathrm{m}} - (b_A + \alpha)I_A^{\mathrm{m}} \qquad (S18d)$$

Both systems may be written in the form:

$$\frac{\mathrm{d}}{\mathrm{d}t}\begin{pmatrix} S_J^{\mathrm{m}} \\ S_A^{\mathrm{m}} \\ I_J^{\mathrm{m}} \\ I_A^{\mathrm{m}} \end{pmatrix} = J\begin{pmatrix} S_J^{\mathrm{m}} \\ S_A^{\mathrm{m}} \\ I_J^{\mathrm{m}} \\ I_A^{\mathrm{m}} \end{pmatrix} \qquad (S19a)$$

where the Jacobian matrix, $J$, is given by:

$$J = \begin{pmatrix} -b_J - g - \lambda_J^* & a(1-N^*) & 0 & af(1-N^*) \\ g & -b_A - \lambda_A^* & 0 & 0 \\ \lambda_J^* & 0 & -b_J - g - \alpha & 0 \\ 0 & \lambda_A^* & g & -b_A - \alpha \end{pmatrix} \qquad (S19b)$$

Note that when $a$, $g$, $b_J$ or $b_A$ trade off with a resistance trait, this is the mutant juvenile resistance and the resident adult resistance in the case of the juvenile resistance mutant invasion dynamics and vice versa in the case of the adult resistance mutant invasion dynamics. $\lambda_J^*$ and $\lambda_A^*$ are similarly functions of the mutant or resident resistance trait depending on whether the juvenile or adult resistance mutant invasion dynamics are being considered. Where the endemic equilibrium ($S_J^*$, $S_A^*$, $I_J^*$, $I_A^*$ and $N^*$) depends on a resistance trait, this is always the resident trait.

We can then calculate the next-generation matrix using the following decomposition of $J$ (which separates out the terms representing new births from the terms which represent deaths or movement between classes):

$$J = \begin{pmatrix} 0 & a(1-N^*) & 0 & af(1-N^*) \\ 0 & 0 & 0 & 0 \\ 0 & 0 & 0 & 0 \\ 0 & 0 & 0 & 0 \end{pmatrix} - \begin{pmatrix} b_J + g + \lambda_J^* & 0 & 0 & 0 \\ -g & b_A + \lambda_A^* & 0 & 0 \\ -\lambda_J^* & 0 & b_J + g + \alpha & 0 \\ 0 & -\lambda_A^* & -g & b_A + \alpha \end{pmatrix} \qquad (S20)$$

This then gives the following next-generation matrix (see [1] for method):

$$N_G = \frac{a(1-N^*)}{D} \begin{pmatrix} N_1 & N_2 & N_3 & N_4 \\ 0 & 0 & 0 & 0 \\ 0 & 0 & 0 & 0 \\ 0 & 0 & 0 & 0 \end{pmatrix} \qquad (S21a)$$

where:

$$D = \left(b_J + g + \lambda_J^*\right)(b_A + \lambda_A^*)\left(b_J + g + \alpha\right)(b_A + \alpha) \qquad (S21b)$$

$$N_1 = g\left(b_J + g + \alpha\right)(b_A + \alpha) + fg\lambda_A^*\left(b_J + g + \alpha\right) + fg\lambda_J^*(b_A + \lambda_A^*) \qquad (S21c)$$

$$N_2 = \left(b_J + g + \lambda_J^*\right)\left(b_J + g + \alpha\right)(b_A + \alpha) + f\lambda_A^*\left(b_J + g + \alpha\right)\left(b_J + g + \lambda_J^*\right) \qquad (S21d)$$

$$N_3 = fg(b_A + \lambda_A^*)\left(b_J + g + \lambda_J^*\right) \qquad (S21e)$$

$$N_4 = f\left(b_J + g + \lambda_J^*\right)(b_A + \lambda_A^*)\left(b_J + g + \alpha\right) \qquad (S21f)$$

In the case where juvenile resistance trades off with maturation and adult resistance trades off with reproduction, the invasion fitness for the juvenile resistance mutant is then:

$$w_J\left(r_J^{\mathrm{m}}, r_J, r_A\right) = \frac{g\left(r_J^{\mathrm{m}}\right)a(r_A)(1-N^*)A_J^{\mathrm{m}}}{(b_A+\alpha)\left(b_J+g\left(r_J^{\mathrm{m}}\right)+\alpha\right)\left(b_A+\lambda_A^*(r_A)\right)\left(b_J+g\left(r_J^{\mathrm{m}}\right)+\lambda_J^*\left(r_J^{\mathrm{m}}\right)\right)} - 1 \qquad (S22a)$$

where, for notational convenience:

$$A_J^{\mathrm{m}} = (b_A+\alpha)\left(b_J+g\left(r_J^{\mathrm{m}}\right)+\alpha\right) + f\left(b_J+g\left(r_J^{\mathrm{m}}\right)+\alpha\right)\lambda_A^*(r_A) + f\lambda_J^*\left(r_J^{\mathrm{m}}\right)\left(b_A+\lambda_A^*(r_A)\right) \qquad (S22b)$$

Similarly, for the adult resistance mutant above, the invasion fitness is:

$$w_A\left(r_A^{\mathrm{m}}, r_J, r_A\right) = \frac{g\left(r_J\right)a(r_A^{\mathrm{m}})(1-N^*)A_A^{\mathrm{m}}}{(b_A+\alpha)\left(b_J+g\left(r_J\right)+\alpha\right)\left(b_A+\lambda_A^*(r_A^{\mathrm{m}})\right)\left(b_J+g\left(r_J\right)+\lambda_J^*\left(r_J\right)\right)} - 1 \qquad (S23a)$$

where, for notational convenience:

$$A_A^{\mathrm{m}} = (b_A+\alpha)\left(b_J+g\left(r_J\right)+\alpha\right) + f\left(b_J+g\left(r_J\right)+\alpha\right)\lambda_A^*(r_A^{\mathrm{m}}) + f\lambda_J^*\left(r_J\right)\left(b_A+\lambda_A^*(r_A^{\mathrm{m}})\right) \qquad (S23b)$$

Similar expressions may be derived for other combinations of trade-offs.

#### Class distribution and reproductive values

Alternatively, we can calculate the next-generation matrix using a different decomposition of $J$ (which separates out the terms representing new additions to each class from the terms which represent removal from a class):

$$J = \begin{pmatrix} 0 & a(1-N^*) & 0 & af(1-N^*) \\ g & 0 & 0 & 0 \\ \lambda_J^* & 0 & 0 & 0 \\ 0 & \lambda_A^* & g & 0 \end{pmatrix} - \begin{pmatrix} b_J+g+\lambda_J^* & 0 & 0 & 0 \\ 0 & b_A+\lambda_A^* & 0 & 0 \\ 0 & 0 & b_J+g+\alpha & 0 \\ 0 & 0 & 0 & b_A+\alpha \end{pmatrix} \qquad (S24)$$

This gives an alternative next-generation matrix:

$$N_G = \frac{1}{D}\begin{pmatrix} 0 & a(1-N^*)v_1v_3v_4 & 0 & af(1-N^*)v_1v_2v_3 \\ gv_2v_3v_4 & 0 & 0 & 0 \\ \lambda_J^*v_2v_3v_4 & 0 & 0 & 0 \\ 0 & \lambda_A^*v_1v_3v_4 & gv_1v_2v_4 & 0 \end{pmatrix} \qquad (S25a)$$

where:

$$D = (b_J + g + \lambda_J^*)(b_A + \lambda_A^*)(b_J + g + \alpha)(b_A + \alpha) \qquad (S25b)$$

$$v_1 = b_J + g + \lambda_J^* \qquad (S25c)$$

$$v_2 = b_A + \lambda_A^* \qquad (S25d)$$

$$v_3 = b_J + g + \alpha \qquad (S25e)$$

$$v_4 = b_A + \alpha \qquad (S25f)$$

Note that this next-generation matrix arises from considering a generation to be the time spent by an individual in a particular class (rather than its entire lifetime). We cannot use this version of the next-generation matrix to find a proxy for the invasion fitness because we cannot determine its eigenvalues analytically.

Let us now consider a monomorphic population (where $r_J^{\mathrm{m}} = r_J$ and $r_A^{\mathrm{m}} = r_A$). In this case, we can calculate the eigenvalues and eigenvectors of the next-generation matrix above.

The right eigenvector is proportional to the class distribution:

$$\begin{pmatrix} (b_J + g + \lambda_J^*)(b_A + \lambda_A^*)(b_J + g + \alpha) \\ g(b_A + \lambda_A^*)(b_J + g + \alpha) \\ \lambda_J^*(b_A + \lambda_A^*)(b_J + g + \alpha) \\ \lambda_J^* g(b_A + \lambda_A^*) + \lambda_A^* g(b_J + g + \alpha) \end{pmatrix} \qquad (S26)$$

The left eigenvector is proportional to the reproductive values (the proportional contribution to distant future generations by individuals currently in each class):

$$R = \begin{pmatrix} g(b_J + g + \alpha)(b_A + \alpha) + fg\lambda_A^*(b_J + g + \alpha) + fg\lambda_J^*(b_A + \lambda_A^*) \\ (b_J + g + \lambda_J^*)(b_J + g + \alpha)(b_A + \alpha) + f\lambda_A^*(b_J + g + \lambda_J^*)(b_J + g + \alpha) \\ fg(b_J + g + \lambda_J^*)(b_A + \lambda_A^*) \\ f(b_J + g + \lambda_J^*)(b_A + \lambda_A^*)(b_J + g + \alpha) \end{pmatrix}^{\mathrm{T}} \qquad (S27)$$

Note that this is also proportional to $(N_1 \quad N_2 \quad N_3 \quad N_4)$, the top row of our original next-generation matrix.

#### FITNESS GRADIENTS

We can differentiate our expressions for the invasion fitness to find the fitness gradients. These can be written in terms of the reproductive values (we will denote the reproductive values given above by $R = (R_1 \quad R_2 \quad R_3 \quad R_4)$). Taking all mutant traits to be evaluated at the resident trait values, we have:

$$\frac{\partial w_J}{\partial r_J^{\mathrm{m}}} = \frac{(1-N^*)}{D}\left(\frac{\partial \lambda_J^*}{\partial r_J^{\mathrm{m}}}\frac{a(R_3-R_1)}{b_J+g+\lambda_J^*} + \frac{\partial a}{\partial r_J^{\mathrm{m}}}R_1 + \frac{\mathrm{d}g}{\mathrm{d}r_J^{\mathrm{m}}}\frac{a(R_2-R_1)(b_J+g+\alpha)+a\lambda_J^*(R_4-R_3)}{(b_J+g+\alpha)(b_J+g+\lambda_J^*)} - \frac{\mathrm{d}b_J}{\mathrm{d}r_J^{\mathrm{m}}}\frac{aR_1(b_J+g+\alpha)+a\lambda_J^*R_3}{(b_J+g+\alpha)(b_J+g+\lambda_J^*)}\right) \quad (S28)$$

$$\frac{\partial w_A}{\partial r_A^{\mathrm{m}}} = \frac{(1-N^*)}{D}\left(\frac{\partial \lambda_A^*}{\partial r_A^{\mathrm{m}}}\frac{ag(R_4-R_2)}{(b_J+g+\lambda_J^*)(b_A+\lambda_A^*)} + \frac{\partial a}{\partial r_A^{\mathrm{m}}}R_1 - \frac{\mathrm{d}b_A}{\mathrm{d}r_A^{\mathrm{m}}}\frac{agR_2(b_J+g+\alpha)(b_A+\alpha)+agR_4\left(\lambda_A^*(b_J+g+\alpha)+\lambda_J^*(b_A+\lambda_A^*)\right)}{D}\right) \quad (S29)$$

#### INTERPRETATION OF FITNESS GRADIENTS

Let us consider the case where the pathogen exerts full sterility virulence ($f = 0$) and no mortality virulence ($\alpha = 0$). Then the fitness gradients can be written as:

$$\frac{\partial w_J}{\partial r_J^{\mathrm{m}}}\Big|_{r_J^{\mathrm{m}}=r_J} = \frac{(1-N^*)(b_J+g)b_A}{D(b_J+g+\lambda_J^*)}\left(-\frac{\partial \lambda_J^*}{\partial r_J}ag + \frac{\partial a}{\partial r_J}g(b_J+g+\lambda_J^*) + \frac{\partial g}{\partial r_J}a(b_J+\lambda_J^*) - \frac{\partial b_J}{\partial r_J}ag\right) \quad (S30a)$$

$$\frac{\partial w_A}{\partial r_A^{\mathrm{m}}}\Big|_{r_A^{\mathrm{m}}=r_A} = \frac{(1-N^*)(b_J+g)b_A}{D(b_A+\lambda_A^*)}\left(-\frac{\partial \lambda_A^*}{\partial r_A}ag + \frac{\partial a}{\partial r_A}g(b_A+\lambda_A^*) - \frac{\partial b_A}{\partial r_A}ag\right) \quad (S30b)$$

These expressions can help us to interpret the effects of different trade-offs. For instance, when both juvenile and adult resistance trade off with reproduction (and if we assume that the natural mortality rates of both age classes are equal), these equations reduce to:

$$\frac{\partial w_J}{\partial r_J} = \frac{(1-N^*)(b_0+g_0)b_0}{D(b_0+g_0+\beta_0(1-r_J)(I_J^*+I_A^*))}\left(\beta_0(I_J^*+I_A^*)a(r_J,r_A)g_0 + \frac{\partial a}{\partial r_J}g_0(b_0+g_0+\beta_0(1-r_J)(I_J^*+I_A^*))\right) \quad (S31a)$$

$$\frac{\partial w_A}{\partial r_A} = \frac{(1-N^*)(b_0+g_0)b_0}{D(b_0+\beta_0(1-r_A)(I_J^*+I_A^*))}\left(\beta_0(I_J^*+I_A^*)a(r_J,r_A)g_0 + \frac{\partial a}{\partial r_A}g_0(b_0+\beta_0(1-r_A)(I_J^*+I_A^*))\right) \quad (S31b)$$

Note that (since $N^* < 1$) the sign of the fitness gradients only depends on the following terms (which appear in the large brackets above):

$$x_J := \beta_0\left(I_J^* + I_A^*\right)a(r_J, r_A)g_0 + \frac{\partial a}{\partial r_J}g_0\left(b_0 + g_0 + \beta_0\left(1 - r_J\right)\left(I_J^* + I_A^*\right)\right) \qquad (S32a)$$

$$x_A := \beta_0\left(I_J^* + I_A^*\right)a(r_J, r_A)g_0 + \frac{\partial a}{\partial r_A}g_0\left(b_0 + \beta_0(1 - r_A)(I_J^* + I_A^*)\right) \qquad (S32b)$$

Co-singular strategies occur for values of the resistance traits at which both terms change sign.

Consider a phase plane with juvenile and adult nullclines which intersect at a co-singular strategy. When juvenile and adult resistance are both sufficiently low, both fitness gradients will be positive (because both traits need to rise to reach the co-singular strategy). Suppose that we move through the phase plane along the line where juvenile and adult resistance are equal. The signs of the fitness gradients along this line are determined by:

$$x_J := \beta_0\left(I_J^* + I_A^*\right)a(r_J, r_J)g_0 + a'g_0\left(b_0 + \beta_0\left(1 - r_J\right)\left(I_J^* + I_A^*\right) + g_0\right) \qquad (S33a)$$

$$x_A := \beta_0\left(I_J^* + I_A^*\right)a(r_J, r_J)g_0 + a'g_0\left(b_0 + \beta_0\left(1 - r_J\right)(I_J^* + I_A^*)\right) \qquad (S33b)$$

where $a' = \frac{\partial a}{\partial r_J}|_{r_A = r_J} = \frac{\partial a}{\partial r_A}|_{r_A = r_J}$ since $r_A = r_J$ on this line.

We can see that $x_J < x_A$ for all values along the line $r_A = r_J$ (since $a' < 0$). Therefore, as we move along the line $r_A = r_J$, the juvenile fitness gradient will change sign from positive to negative before the adult fitness gradient changes sign (it will still be positive at this point). Note that such a sign change must occur since, for sufficiently large values of the juvenile and adult resistance, both fitness gradients must be negative.

Having reached the juvenile nullcline, we can then move along it, away from the line $r_A = r_J$, in the direction where the adult fitness gradient is decreasing (adult resistance increasing). As such, we must be in the $r_A > r_J$ half of the phase plane when we finally reach the co-singular strategy, so we would expect the adult resistance to exceed the juvenile resistance at this point (see the point where $1 - f = 1$ in Fig. 2C).

Note that the difference between $x_J$ and $x_A$ in equations (S33) above is based on the fact that $b_0 + \lambda_J^* + g_0 > b_0 + \lambda_A^*$. These terms represent the rate at which hosts leave the susceptible juvenile and susceptible adult classes respectively and so this inequality shows that, on average, hosts spend more time as susceptible adults than as susceptible juveniles (resistance traits and natural mortality rates being equal). This explains why there is greater selection for adult resistance than juvenile resistance in this case. We can also see that the difference between $x_J$ and $x_A$ depends on the maturation rate, $g_0$. As such, we would expect the difference between juvenile and adult resistance to increase for faster maturation rates, which is indeed what we see in Fig. S10C. Biologically, one can intuit this by taking it to an extreme: if individuals mature immediately after they are born, then there is no advantage to being resistant as a juvenile because the duration of protection is so short.

Similar arguments can be applied to interpret other combinations of trade-offs (still with $f = 0, \alpha = 0$ for tractability). For instance, consider the same juvenile trade-off but now with an adult resistance/mortality trade-off. We have $\left|\frac{1}{a_0}\frac{\partial a}{\partial r_J}\right| a_0 g_0 (b_0 + g_0 + \lambda_J^*) > \left|\frac{1}{b_0}\frac{\partial b_A}{\partial r_A}\right| b_0 a g_0$ along the line $r_A = r_J$ since $a \le a_0$ and the trade-off gradients, when divided by their scaling constants, are of the same magnitude. Hence, the juvenile fitness gradient passes through zero first, whilst the adult fitness gradient remains positive and so juvenile resistance evolves to be lower than adult resistance, as shown in Fig. 2F. The same argument applies when juvenile resistance trades off with maturation and adult resistance trades off with mortality (Fig. 2D), since we know that $\left|\frac{1}{g_0}\frac{\partial g}{\partial r_J}\right| a_0 g_0 (b_0 + \lambda_J^*) > \left|\frac{1}{b_0}\frac{\partial b_A}{\partial r_A}\right| b_0 a_0 g$. In the case of a juvenile resistance/maturation trade-off and adult resistance/reproduction trade-off, the relevant terms in equation (S30a-b) are equal when the mortality rates are the same, which is why the two resistance traits evolve to the same level at maximal sterility virulence ($f = 0$) in Fig. 2A. The same argument also applies when both resistance traits trade-off with mortality (Fig. 2E). In the case of juvenile resistance/mortality and adult resistance/reproduction trade-offs, the relevant terms in equation (S30a-b) show that the juvenile term is less than the adult term, $\left|\frac{1}{b_0}\frac{\partial b_J}{\partial r_J}\right| b_0 a g_0 < \left|\frac{1}{a_0}\frac{\partial a}{\partial r_A}\right| a_0 g_0 (b_0 + \lambda_A^*)$, implying that juvenile resistance will evolve to be higher than adult resistance (Fig. 2B).

#### STABILITY CONDITIONS

A co-singular strategy $(r_J^*,\ r_A^*)$ is co-evolutionarily stable if and only if $\frac{\partial^2 w_J}{\partial r_J^{\mathrm{m}2}} < 0$ and $\frac{\partial^2 w_A}{\partial r_A^{\mathrm{m}2}} < 0$ at the co-singular strategy [2].

A co-singular strategy is strong convergence stable if and only if $\frac{\partial^2 w_J}{\partial r_J^{\mathrm{m}2}} + \frac{\partial^2 w_J}{\partial r_J^{\mathrm{m}} \partial r_J} < 0$ and $\frac{\partial^2 w_A}{\partial r_A^{\mathrm{m}2}} + \frac{\partial^2 w_A}{\partial r_A^{\mathrm{m}} \partial r_A} < 0$ and $\frac{\partial^2 w_J}{\partial r_J^{\mathrm{m}} \partial r_A}\frac{\partial^2 w_A}{\partial r_A^{\mathrm{m}} \partial r_J} - \left(\frac{\partial^2 w_J}{\partial r_J^{\mathrm{m}2}} + \frac{\partial^2 w_J}{\partial r_J^{\mathrm{m}} \partial r_J}\right)\left(\frac{\partial^2 w_A}{\partial r_A^{\mathrm{m}2}} + \frac{\partial^2 w_A}{\partial r_A^{\mathrm{m}} \partial r_A}\right) < 0$ at the co-singular strategy [2].

Note that this method assumes that the juvenile and adult resistance traits have equal mutation rates.

#### DESCRIPTION OF NUMERICAL METHODS (EVOLUTIONARY INVASION ANALYSIS)

1. Create matrices of the juvenile and adult fitness gradients evaluated at different values of juvenile and adult resistance ranging from zero to one.
2. Determine values of juvenile and adult resistance at which both matrices change sign. These are the co-singular strategies.
3. If there are no sign changes in either matrix then the co-singular strategy will comprise juvenile resistance at zero (if the juvenile fitness gradient matrix is always negative) or one (if the juvenile matrix is always positive) and similarly for adult resistance. If there are sign changes then repeat steps 1 and 2 for values of juvenile and adult resistance over a smaller range near to the sign change (to give the co-singular strategy more accurately).
4. Record the co-singular strategies.
5. If there are many singular strategies close together due to numerical errors then organise these into 'clusters' of singular strategies with very similar values. Use an ODE solver to find roots of the fitness gradients over the range of values of juvenile and adult

resistance present in the cluster. If a singular strategy is found then replace the cluster by this. If no singular strategy is found then replace the cluster by an average value of juvenile and adult resistance over the cluster and display a warning message. If there are no clusters of singular strategies close together then retain the original list of co-singular strategies.

6. Calculate the ecological, endemic equilibrium at each co-singular strategy.
7. Substitute the values of juvenile and adult resistance at each co-singular strategy and the ecological equilibrium into expressions for the second derivatives of the invasion fitness.
8. Use the second derivatives of the invasion fitness to determine the evolutionary stability and strong convergence stability of the co-singular strategies.

This method assumes that mutations are rare (allowing separation of ecological and evolutionary timescales) and that mutations have small phenotypic effects. We relax these assumptions using evolutionary simulations, thereby verifying our results.

##### DESCRIPTION OF EVOLUTIONARY SIMULATIONS

1. Run the ecological dynamics of the system for a fixed length of time, with the juvenile and adult resistance taking their resident values.
2. Introduce a mutant, randomly determining whether the mutation will occur in the juvenile or adult resistance (we assume that both resistance traits evolve at the same rate) and whether the resistance trait will mutate to be slightly higher or lower than its current value. Add a small sub-population with the new, mutant trait values.
3. Run the ecological dynamics of the system for a fixed length of time, starting at its current composition.
4. Remove any sub-populations which have a density below a low threshold (they are extinct).
5. Introduce a mutant by randomly determining which sub-population (combination of juvenile and adult resistance trait values) the mutant will come from, whether it will mutate in the juvenile or adult resistance trait and whether the resistance trait will mutate to be slightly higher or lower than its current value. Add a small sub-population with the new, mutant trait values.

6. Repeat steps 3 to 5 for many evolutionary timesteps.

In these simulations, ecological and evolutionary timescales are not completely separated because the ecological system does not necessarily reach its equilibrium before the next mutant is introduced. Mutations do not have arbitrarily small phenotypic effects because mutations alter resistance trait values by a small, fixed interval.

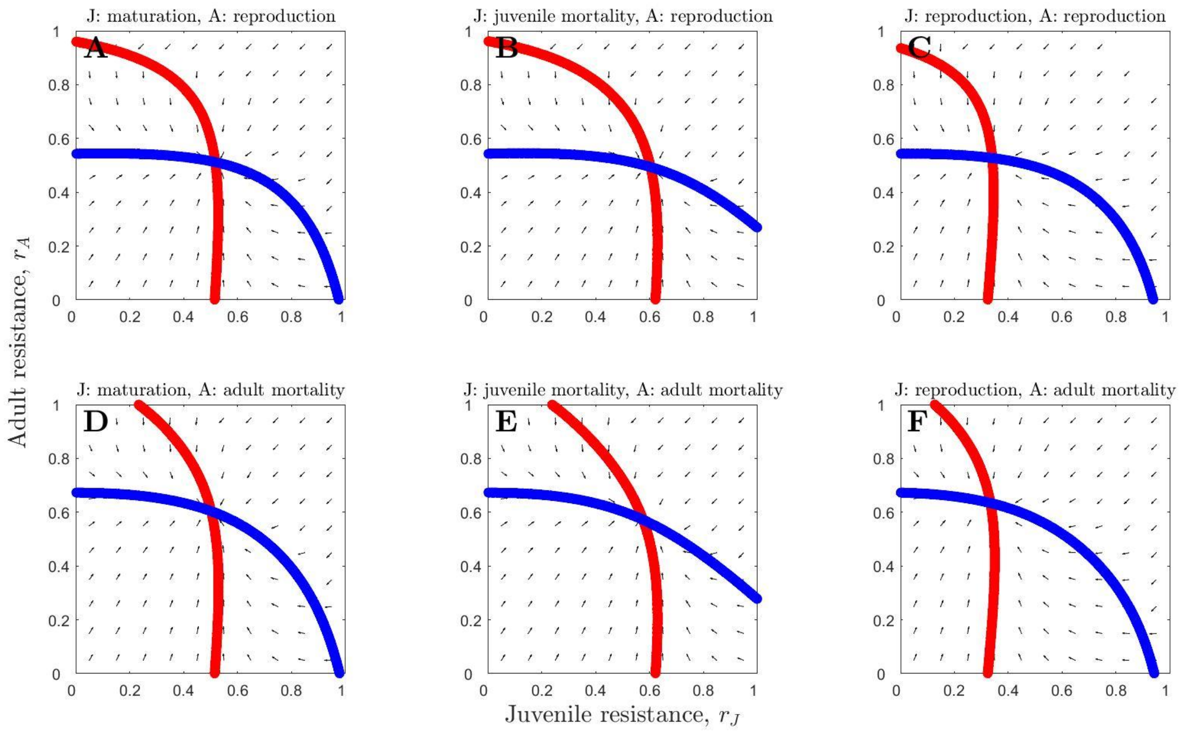


Fig. S1: Phase planes showing continuously stable strategies, with the juvenile nullcline in red and the adult nullcline in blue, for six different combinations of trade-offs: (A)-(C) adult resistance with reproduction, (D)-(F) adult resistance with adult mortality, (A) & (D) juvenile resistance with maturation, (B) & (D) juvenile resistance with juvenile mortality and (C) & (F) juvenile resistance with reproduction. We can see that the juvenile nullcline (red) is shifted significantly to the left when juvenile resistance trades off with reproduction (C and F) as opposed to maturation or mortality (A-B, D-E). Parameter values are as in Table 1, with $\beta_0 = 8$, $\alpha = 0$ and $f = 0.1$.

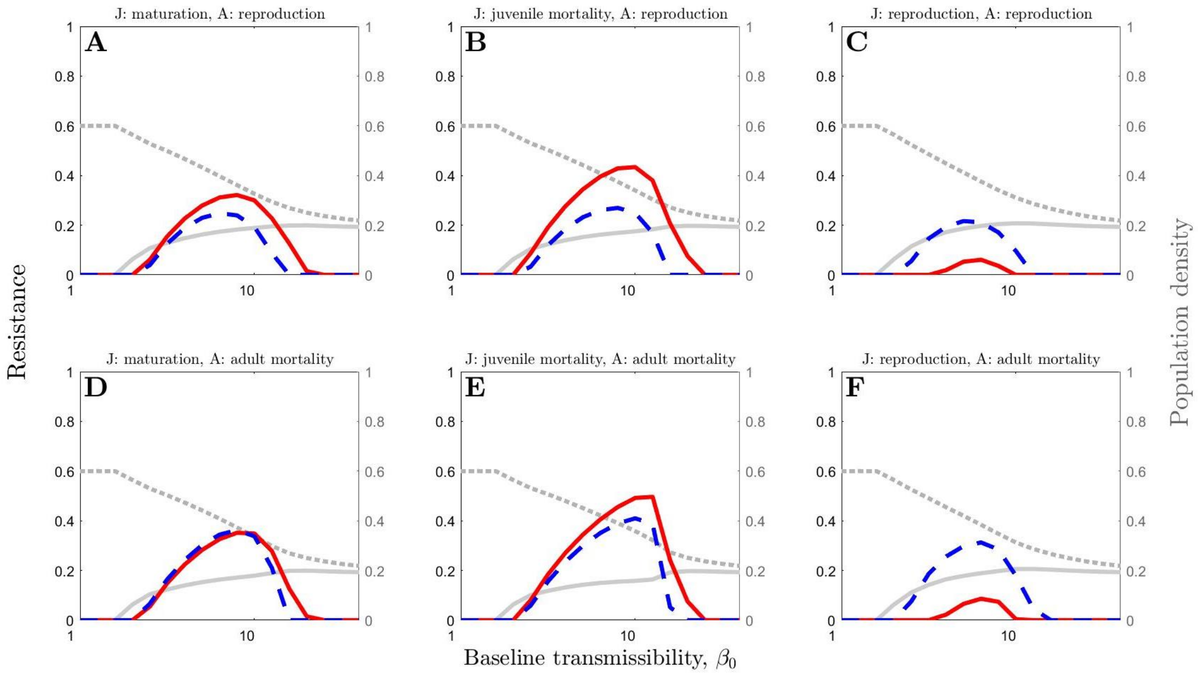


Fig. S2: The effect of varying baseline transmissibility, $\beta_0$, on juvenile resistance (solid red) and adult resistance (dashed blue), for six different combinations of trade-offs. The dotted, grey line shows total population density and the solid, grey line shows the density of infected hosts (both are non-dimensionalised). Parameter values are as in Table 1, with $\alpha = 0$ and $f = 0.5$.

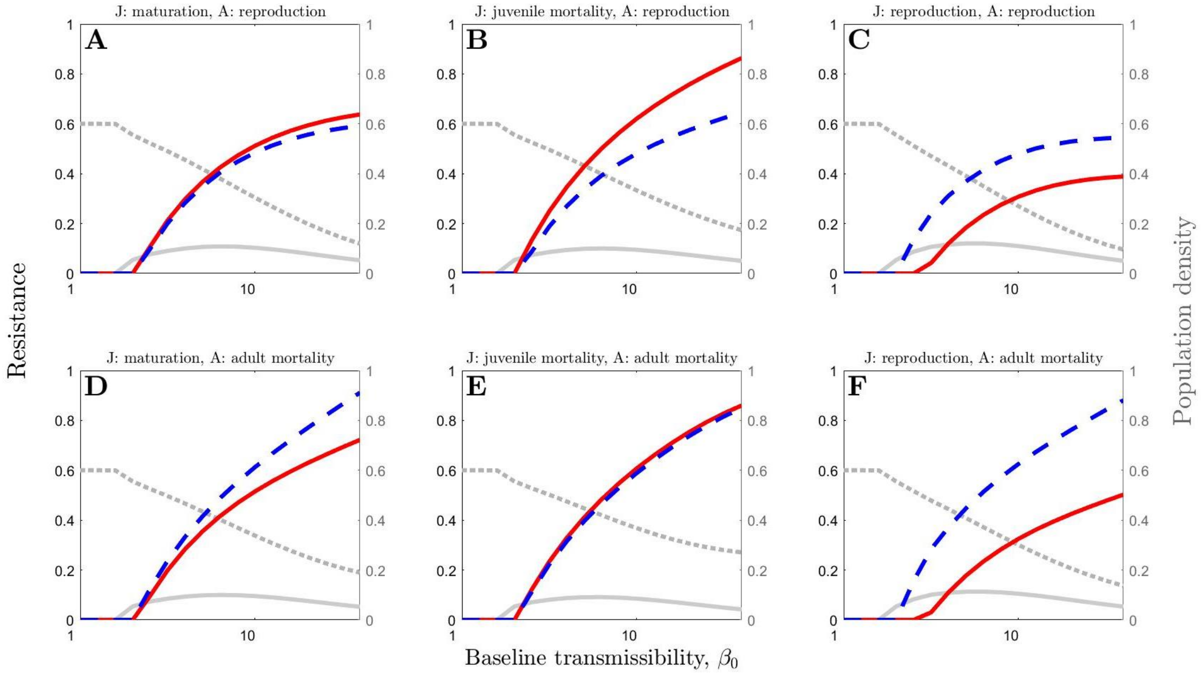


Fig. S3: The effect of varying baseline transmissibility, $\beta_0$, on juvenile resistance (solid red) and adult resistance (dashed blue), for six different combinations of trade-offs. The dotted, grey line shows total population density and the solid, grey line shows the density of infected hosts (both are non-dimensionalised). Parameter values are as in Table 1, with $\alpha = 0$ and $f = 0.3$.

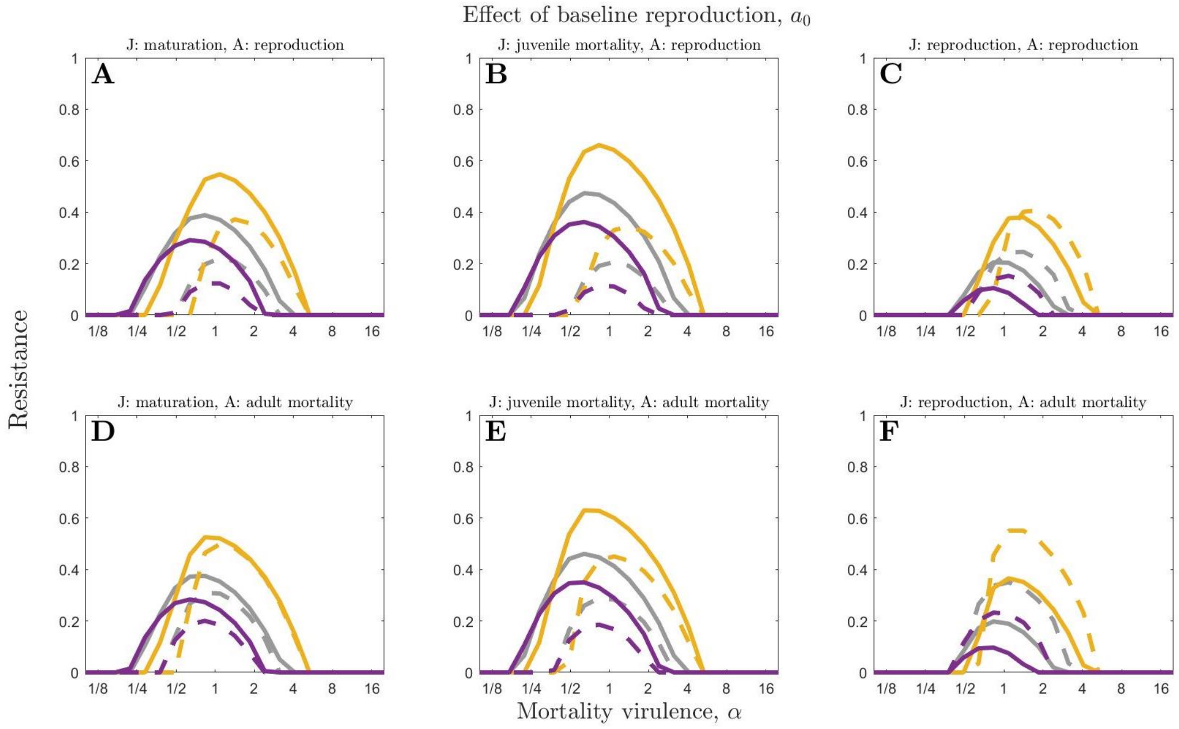


Fig. S4: The effect of varying mortality virulence, $\alpha$, on juvenile resistance (solid) and adult resistance (dashed), for three different values of baseline reproduction rate: a high value ($a_0 = 10$; orange), an intermediate value ($a_0 = 5$; grey) and a low value ($a_0 = 4$; purple). Results are shown for six different combinations of trade-offs. Parameter values are as in Table 1, with $\beta_0 = 8$ and $f = 1$.

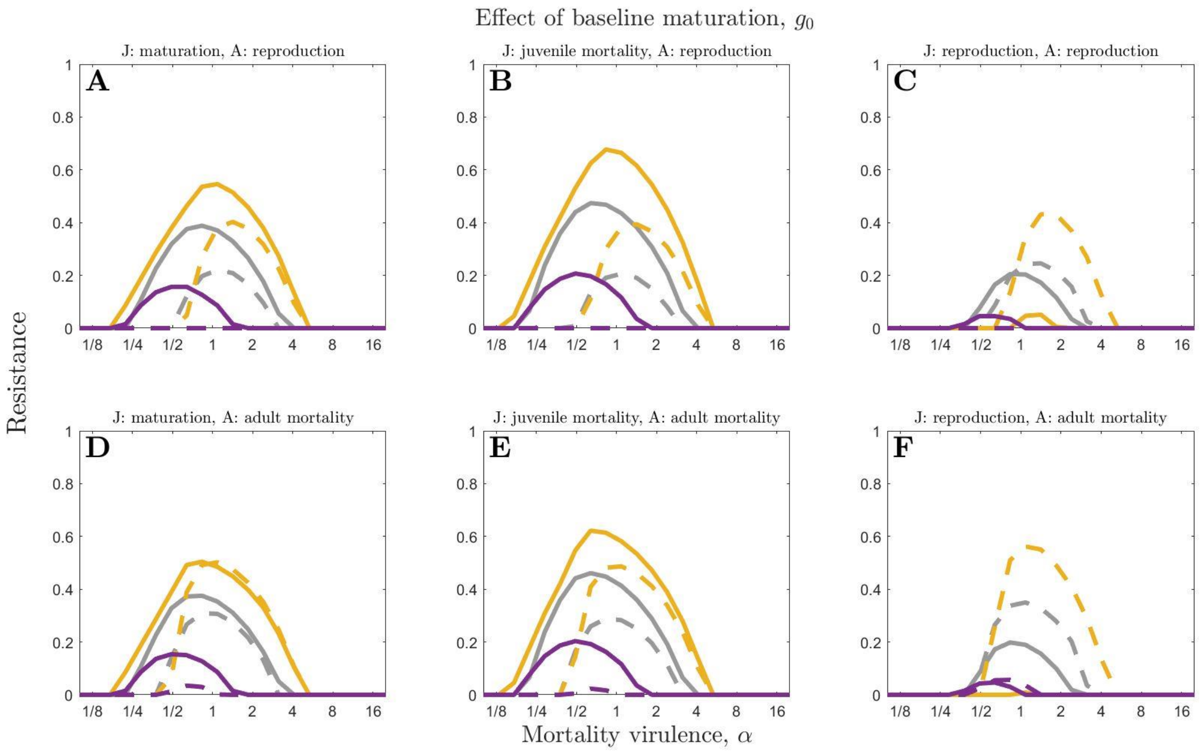


Fig. S5: The effect of varying mortality virulence, $\alpha$, on juvenile resistance (solid) and adult resistance (dashed), for three different values of baseline maturation rate: a high value ($g_0 = 5$; orange), an intermediate value ($g_0 = 1$; grey) and a low value ($g_0 = 0.5$; purple). Results are shown for six different combinations of trade-offs. Parameter values are as in Table 1, with $\beta_0 = 8$ and $f = 1$.

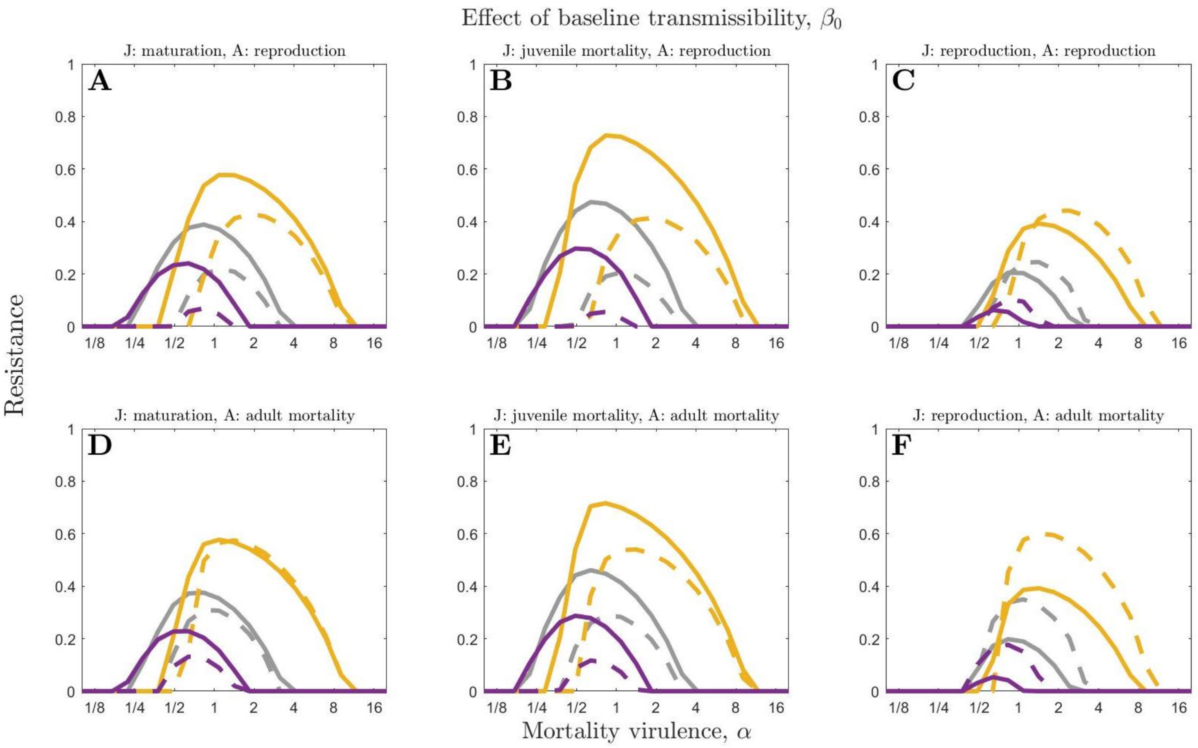


Fig. S6: The effect of varying mortality virulence, $\alpha$, on juvenile resistance (solid) and adult resistance (dashed), for three different values of baseline transmissibility: a high value ($\beta_0 = 20$; orange), an intermediate value ($\beta_0 = 8$; grey) and a low value ($\beta_0 = 5$; purple). Results are shown for six different combinations of trade-offs. Parameter values are as in Table 1, with $f = 1$.

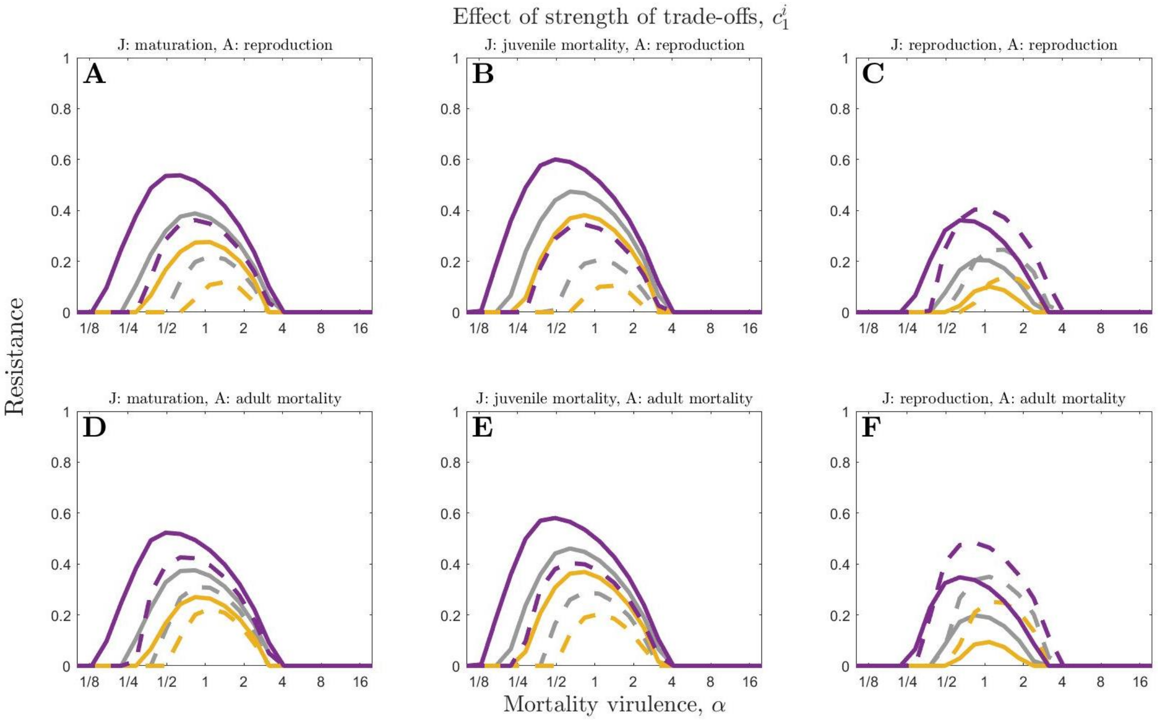


Fig. S7: The effect of varying mortality virulence, $\alpha$, on juvenile resistance (solid) and adult resistance (dashed), for three different values of the strength of the trade-offs: a high value ($c_1^{\mathrm{i}} = 0.7$; orange), an intermediate value ($c_1^{\mathrm{i}} = 0.5$; grey) and a low value ($c_1^{\mathrm{i}} = 0.3$; purple). Results are shown for six different combinations of trade-offs. Parameter values are as in Table 1, with $\beta_0 = 8$ and $f = 1$.

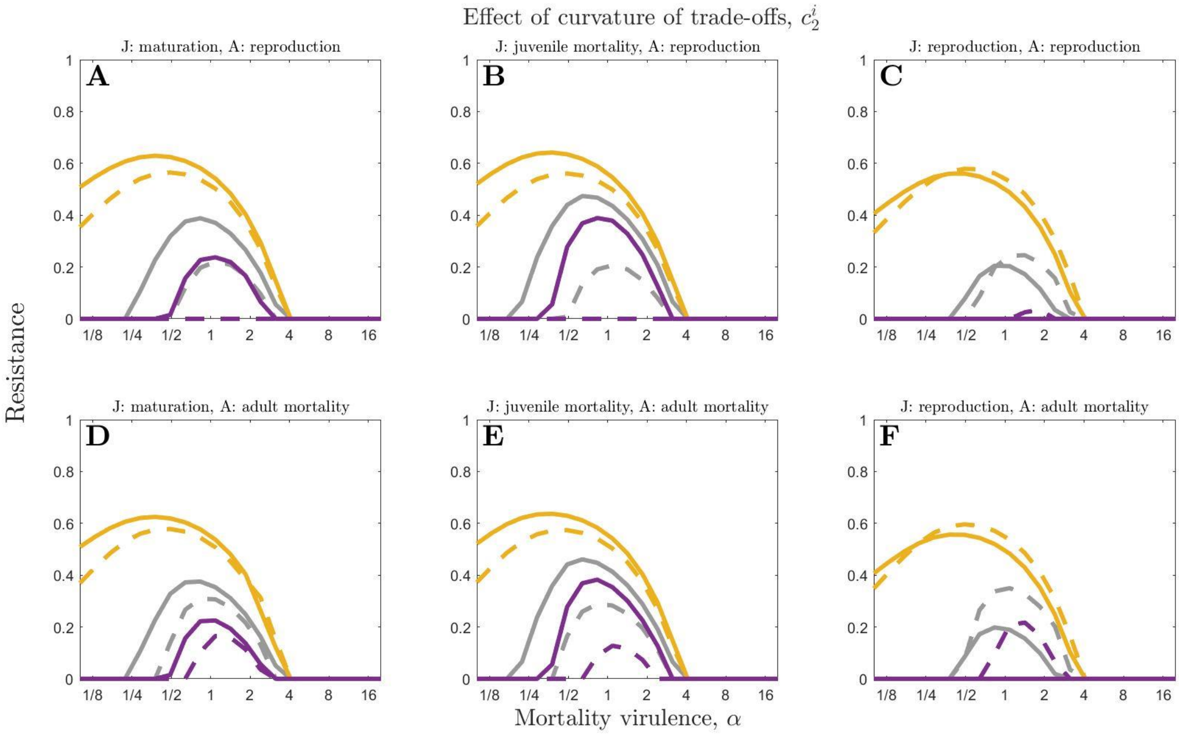


Fig. S8: The effect of varying mortality virulence, $\alpha$, on juvenile resistance (solid) and adult resistance (dashed), for three different values of the curvature of the trade-offs: a high value ($c_2^{\mathrm{i}} = 8$; orange), an intermediate value ($c_2^{\mathrm{i}} = 3$; grey) and a low value ($c_2^{\mathrm{i}} = 2$; purple). Results are shown for six different combinations of trade-offs. Parameter values are as in Table 1, with $\beta_0 = 8$ and $f = 1$.

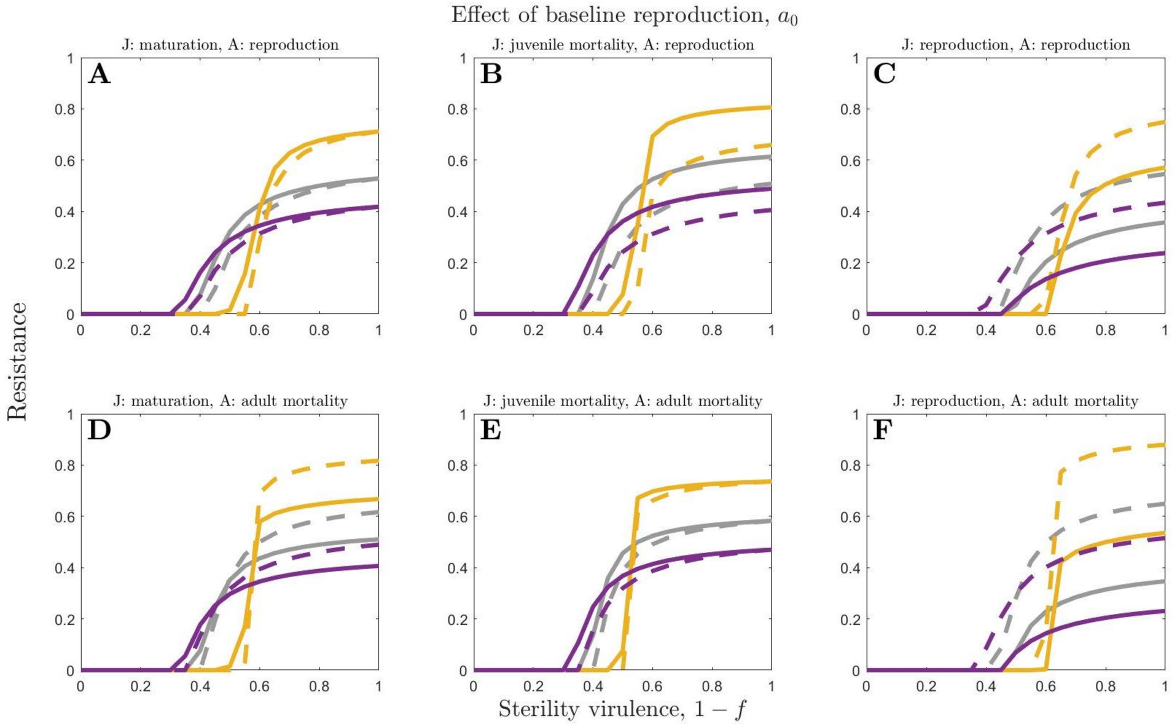


Fig. S9: The effect of varying sterility virulence, $1 - f$, on juvenile resistance (solid) and adult resistance (dashed), for three different values of baseline reproduction rate: a high value ($a_0 = 10$; orange), an intermediate value ($a_0 = 5$; grey) and a low value ($a_0 = 4$; purple). Results are shown for six different combinations of trade-offs. Parameter values are as in Table 1, with $\beta_0 = 8$ and $\alpha = 0$.

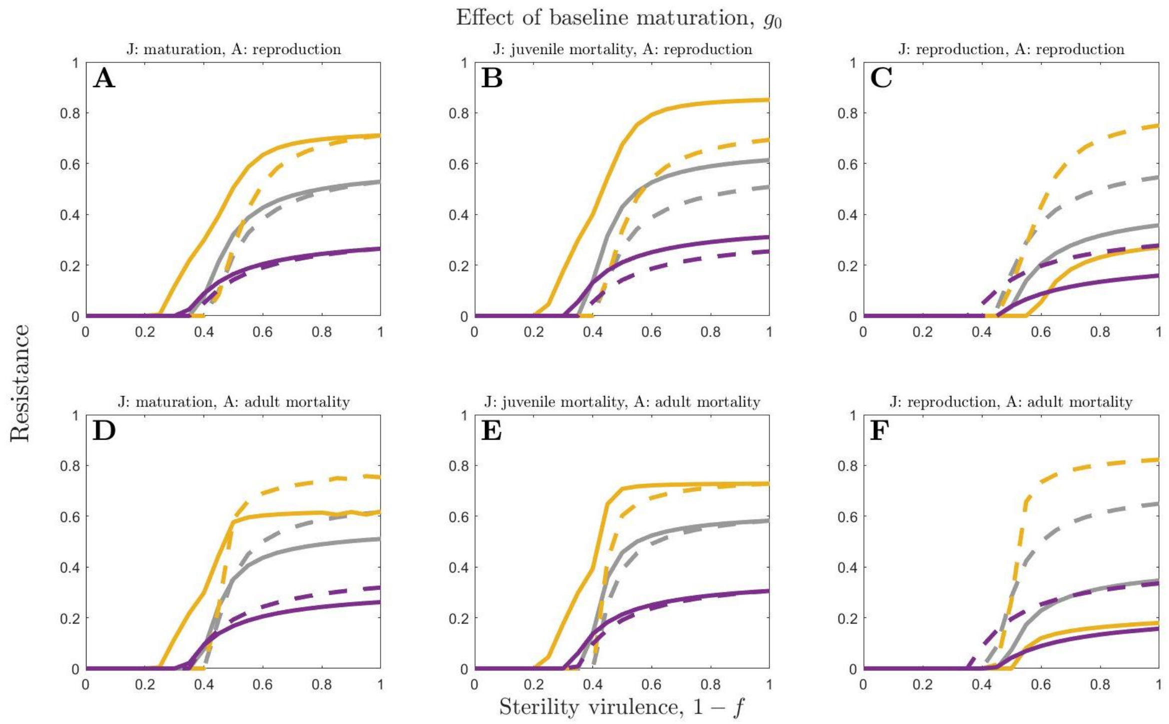


Fig. S10: The effect of varying sterility virulence, $1 - f$, on juvenile resistance (solid) and adult resistance (dashed), for three different values of baseline maturation rate: a high value ($g_0 = 5$; orange), an intermediate value ($g_0 = 1$; grey) and a low value ($g_0 = 0.5$; purple). Results are shown for six different combinations of trade-offs. Parameter values are as in Table 1, with $\beta_0 = 8$ and $\alpha = 0$.

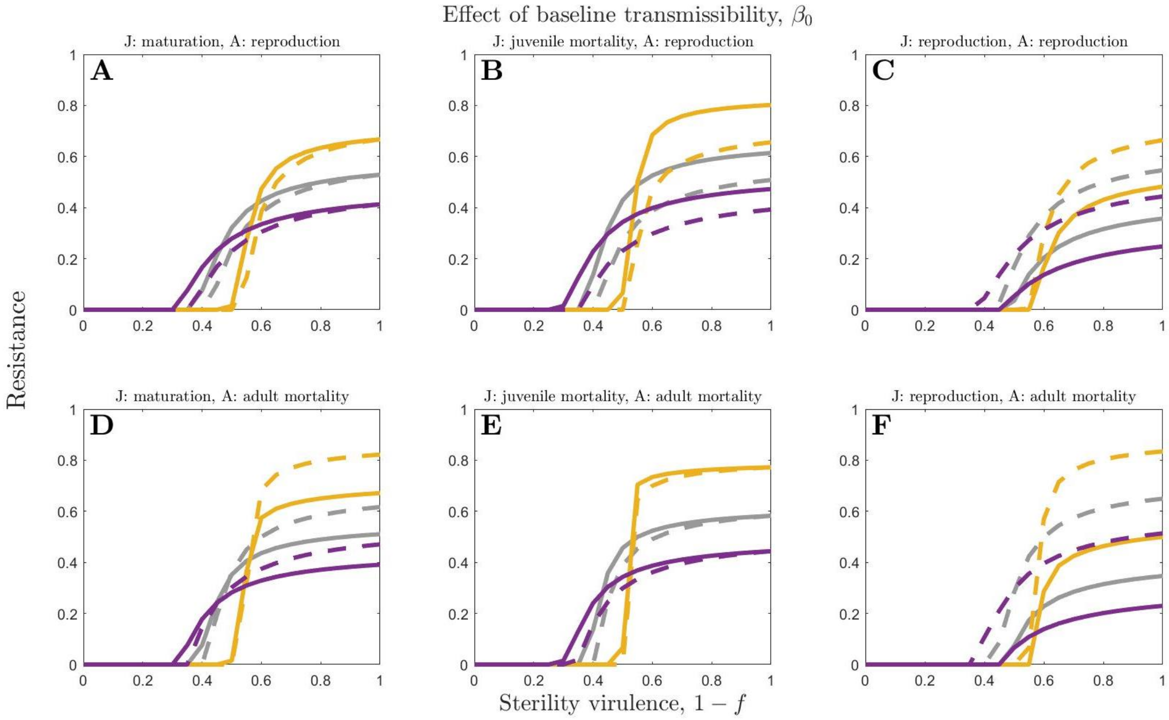


Fig. S11: The effect of varying sterility virulence, $1 - f$, on juvenile resistance (solid) and adult resistance (dashed), for three different values of baseline transmissibility: a high value ($\beta_0 = 20$; orange), an intermediate value ($\beta_0 = 8$; grey) and a low value ($\beta_0 = 5$; purple). Results are shown for six different combinations of trade-offs. Parameter values are as in Table 1, with $\alpha = 0$.

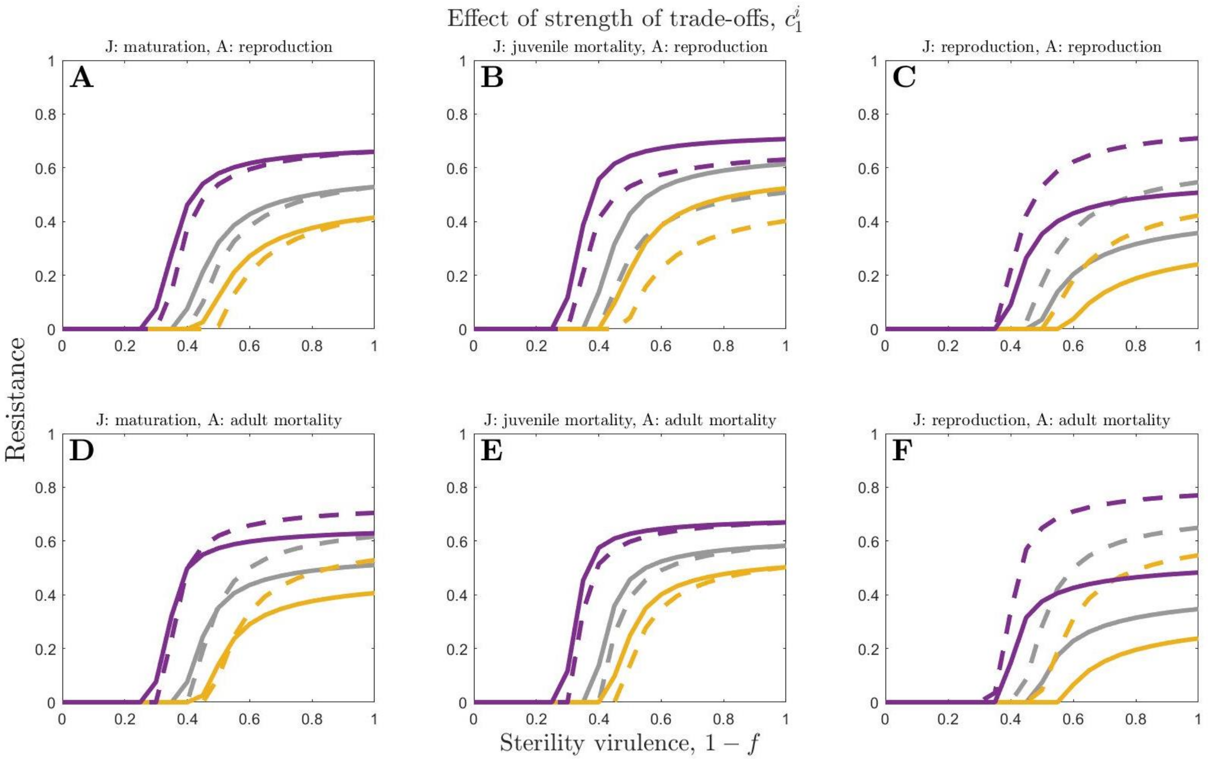


Fig. S12: The effect of varying sterility virulence, $1-f$, on juvenile resistance (solid) and adult resistance (dashed), for three different values of the strength of the trade-offs: a high value ($c_1^{\mathrm{i}} = 0.7$; orange), an intermediate value ($c_1^{\mathrm{i}} = 0.5$; grey) and a low value ($c_1^{\mathrm{i}} = 0.3$; purple). Results are shown for six different combinations of trade-offs. Parameter values are as in Table 1, with $\beta_0 = 8$ and $\alpha = 0$.

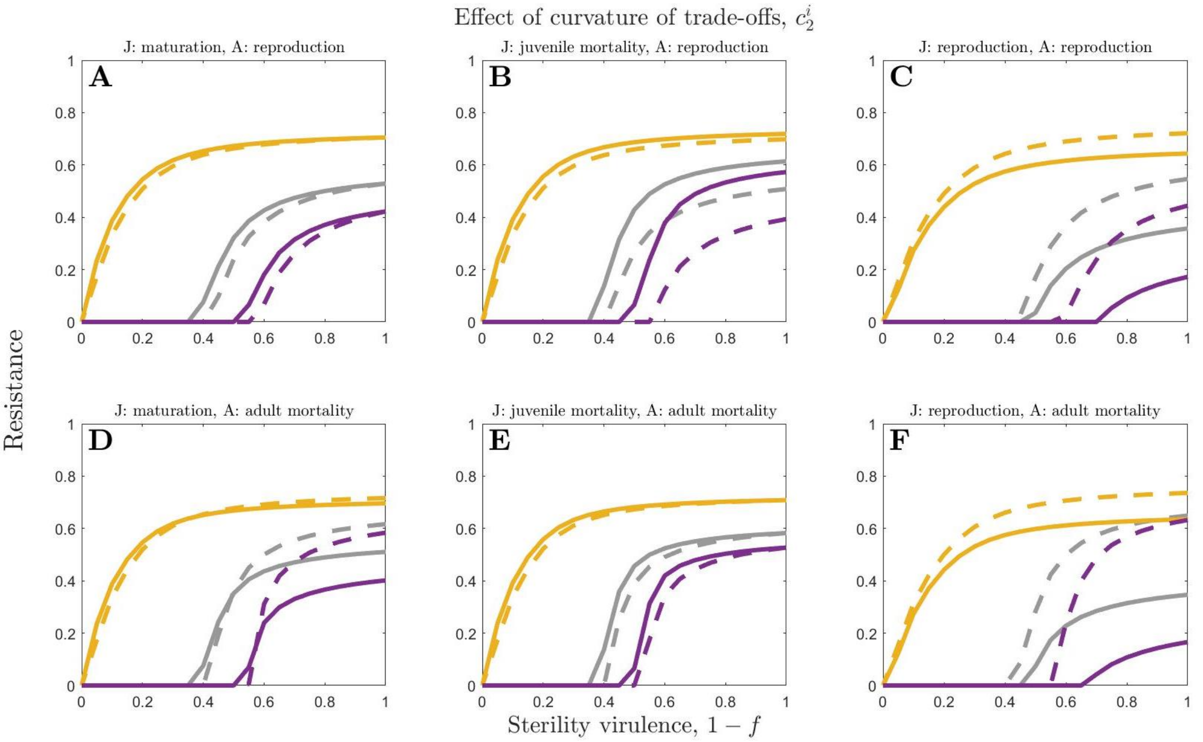


Fig. S13: The effect of varying sterility virulence, $1-f$, on juvenile resistance (solid) and adult resistance (dashed), for three different values of the curvature of the trade-offs: a high value ($c_2^{\mathrm{i}} = 8$; orange), an intermediate value ($c_2^{\mathrm{i}} = 3$; grey) and a low value ($c_2^{\mathrm{i}} = 2$; purple). Results are shown for six different combinations of trade-offs. Parameter values are as in Table 1, with $\beta_0 = 8$ and $\alpha = 0$.
